# Cooperative siderophore use stabilizes a protective leaf microbiome

**DOI:** 10.64898/2026.03.18.712463

**Authors:** Paolo Stincone, Lukas M. Braun, Caner Bağcı, Marcelo Navarro-Diaz, Alicia I. Pérez-Lorente, Shane P. Farrell, Daniel Gómez-Pérez, Johanna Bode, Karoline Steuer-Lodd, Maryam Mahmoudi, Vasvi Chaudhry, Diego Romero, Allegra T. Aron, Nadine Ziemert, Carlos Molina-Santiago, Eric M. Kemen, Daniel Petras

## Abstract

Plant-associated microbial communities provide crucial protection against pathogens. Specialized metabolites play key roles in plant-microbe and microbe-microbe interactions and, ultimately, in plant health; however, the molecular mechanisms underlying their plant-protecting properties remain largely unknown. Nutrient deficiency (e.g., iron) on leaf surfaces creates intense competition among microbes, driving both antagonism and cooperation. Using a gnotobiotic *Arabidopsis thaliana* model and a synthetic leaf microbial community, we show that community stability and plant protection depend on cooperative siderophore exchange between the basidiomycete yeast *Rhodotorula kratochvilovae* and commensal *Pseudomonas* species. Removal of *Pseudomonas* caused a strong shift in the community metabolome and accumulation of the yeast siderophore rhodotorulic acid (RA). RA selectively promoted the growth of commensal *Pseudomonas* via TonB-dependent transporters, which are absent in pathogenic *Pseudomonas* strains. Inactivation of these transporter genes abolished RA uptake, destabilized the synthetic community, and eliminated protection against *Pseudomonas syringae* infection. RA and *Rhodotorula* also induced host iron-deficiency and jasmonate-related defense metabolites, linking microbial cooperation to plant stress responses. These findings reveal that microbial siderophore exchange acts as a key mechanism that maintains stability in the phyllosphere microbiome. Rather than solely promoting competition, iron-binding compounds can serve as cooperative currencies that align microbial fitness with host protection.

---

Plant microbiomes are fundamental to the health and functioning of terrestrial ecosystems globally, regulating nutrient cycling, bolstering plant resilience, structuring biological communities, and suppressing pathogens, thereby enhancing ecosystem stability ^1,2^. In applied contexts such as habitat restoration, climate adaptation, and agriculture, the integrity of plant–microbiome interactions is critical for maintaining ecosystem services that humans depend on, including soil fertility, water retention, carbon sequestration, and sustainable food production ^3,4^. Given the importance of plant-associated microbiomes to both ecological functioning and human well-being, unravelling their dynamics is essential for predicting ecosystem responses to environmental change and for informing conservation, management, and land-use strategies.

Yet, plant-associated microbiomes worldwide are threatened ^5^ by numerous stressors, including habitat degradation, pollution, and extreme climate ^6,7^, which shift the balance from mutualistic microbes to opportunistic pathogens ^8^. Understanding how these complex microbial communities maintain stability and protect their host under stress is becoming increasingly urgent. The phyllosphere, the aerial parts of plants, represents one of the largest microbial habitats on Earth ^9^, yet the mechanisms governing host-microbe and microbe-microbe interactions are poorly understood. Emerging evidence indicates that certain phyllosphere residents, including commensal *Pseudomonas species*, can suppress plant pathogens ^9,10^ however, the specific molecular and ecological mechanisms mediating this protection remain unclear.

One likely driver is microbial competition for scarce resources such as iron, a micronutrient in minimal supply on leaf surfaces^11^. To survive in iron-depleted environments, many microbes secrete siderophores, small, high-affinity iron-chelating compounds ^12^, that not only promote individual fitness but also mediate competitive and cooperative interactions among community members. These molecules can strongly influence microbial assembly and function ^13^.

Here, we investigated how siderophore-producing commensal microbes and siderophore uptake by the commensal *Pseudomonas* keystone species contribute to the protection of *Arabidopsis thaliana*. Prior work investigating the microbiome of natural *Arabidopsis thaliana* populations ^14^ identified a known yet understudied yeast of the genus *Rhodotorula* as a keystone taxon within the phyllosphere, revealing predominantly positive associations with other microbial members, particularly commensal *Pseudomonas* strains (**Fig. 1a-b**). Using a sterile *Arabidopsis thaliana* model and a representative Synthetic Community (SynCom) composed of core phyllosphere microbial members, we combined dropout co-culture experiments, functional metabolomics, amplicon sequencing, pangenome analyses, gene knockouts, and plant phenotyping to dissect the mechanisms underlying community-level iron acquisition. Our data revealed how siderophore sharing and the presence or absence of TonB-dependent transporters shape microbial competitive dynamics, community stability, and plant protection. Our results provide new insights into the molecular and ecological processes that stabilize beneficial plant–microbiome associations and may have broader implications for microbiome stability in environmental and other host-associated systems, including those relevant to human health.

**Fig. 1.**
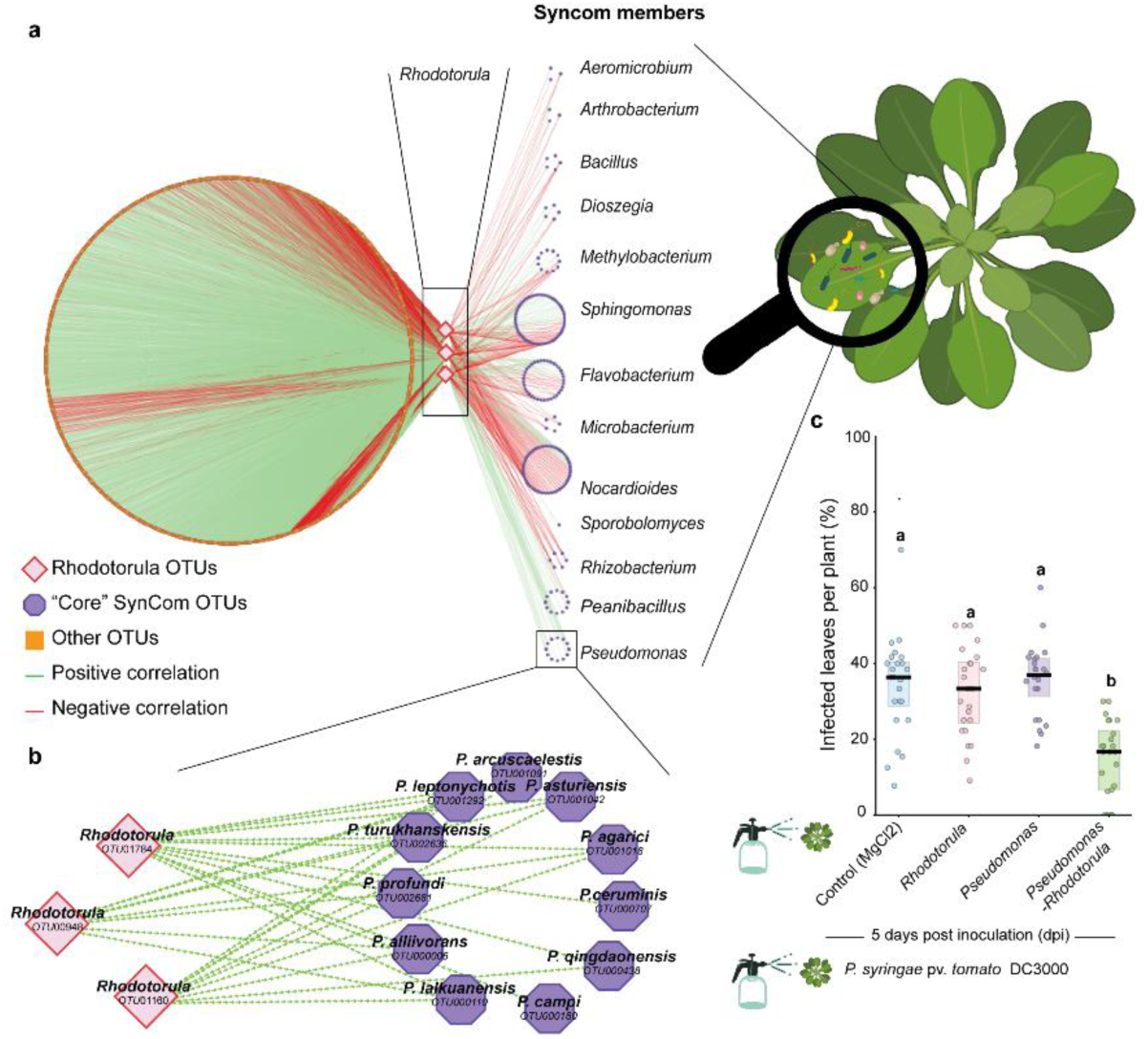
An inter-kingdom positive association in the *Arabidopsis thaliana* phyllosphere protects against the phytopathogen *P. syringae*. **a,** The network displays correlations among *Rhodotorula* Operational Taxonomic Units (OTUs) and other members of the microbial community, highlighting primary correlations represented by the edges. The network encompasses interactions with epiphytic and endophytic microbial communities associated with *Arabidopsis thaliana*. Microbial community data used to construct the network were obtained using amplicon sequencing of the 16S rRNA gene for bacteria, the ITS2 gene for fungi, and the 18S rRNA for eukaryotes other than fungi. Data were collected from six distinct sites around Tübingen, Germany, over five consecutive plant generations. In the visualization, core OTUs are highlighted in violet, and non-core OTUs are displayed in orange, representing taxa considered in other studies. **b,** A detailed view of the inset, highlighting the correlation between *Pseudomonas* and *Rhodotorula* OTUs. Green edges indicate positive correlations between nodes, suggesting co-occurrence or mutualistic interactions. Red edges denote negative correlations, implying antagonistic relationships or competitive interactions. **c,** Percentage of infected leaves per plant following *Pseudomonas syringae* pv. *tomato* DC3000 infection (24 plants per treatment; all leaves scored). One-way ANOVA (*P* = 2.2 × 10⁻⁹) followed by Tukey’s HSD test; different letters indicate significant differences (*P* < 0.05).

## Results

### *Rhodotorula* promotes the growth of commensal *Pseudomonas* and contributes to plant protection

Field surveys conducted across six locations around Tübingen, Germany, over five years revealed strong positive correlations between *Rhodotorula* Operational Taxonomic Units (OTUs) and multiple bacterial taxa, including members of the *Pseudomonas* clade (**Fig. 1a-b**). These patterns suggested possible inter-kingdom facilitation or metabolic complementarity rather than antagonism within the microbiome. To validate these ecological associations experimentally, we used a previously established 15-member SynCom containing representative bacterial and yeast strains from the natural phyllosphere ^15^. We quantified *in vitro* growth responses of all 15 SynCom members when exposed to the cell-free filtrate from *Rhodotorula kratochvilovae* cultures. Growth responses varied among SynCom members, ranging from stimulation to inhibition or no measurable change, confirming strain-specific sensitivity to *Rhodotorula* exometabolites.

Notably, *Rhodotorula* supernatant significantly enhanced the growth of the *P. koreensis* strain present in the SynCom (P<0.05); (**fig. S1**). This *in vitro* growth promotion mirrors the strong co-abundance patterns observed in natural *Arabidopsis* populations (**Fig. 1a-b**), indicating that *Rhodotorula* may directly facilitate the establishment of beneficial *Pseudomonas* lineages in the phyllosphere.

Because commensal *Pseudomonas* are known antagonists of phytopathogens ^9,10^, including *Pseudomonas syringae*, we tested whether such interactions influenced plant protection. Greenhouse experiments showed that non-sterile plants pre-inoculated with both SynCom members *Pseudomonas* and *Rhodotorula* exhibited significantly reduced disease symptoms upon exposure to *P. syringae* pv. *tomato* DC3000 compared to control treatments (ANOVA with Tukey HSD, *P* = 2.2 × 10⁻⁹; **Fig. 1c).** These results demonstrate that the *Rhodotorula*–*Pseudomonas* interaction not only shapes microbial growth dynamics but also contributes to enhanced host protection.

### *Pseudomonas* removal rewires the community metabolome and triggers rhodotorulic acid accumulation

To quantify the contribution of individual community members to the core phyllosphere microbiome of *Arabidopsis thaliana*, we generated a series of dropout SynComs, each lacking one strain (Δ) from the initial 15-member consortium, which was used as a control (**Fig. 2a**). Amplicon sequencing and non-targeted metabolomics via liquid chromatography tandem mass spectrometry (LC–MS/MS) were used to profile both community composition and metabolic output. When *Pseudomonas koreensis* was omitted (Δ*Pseudomonas*), bacterial genera such as *Bacillus*, *Microbacterium*, and *Methylobacterium* expanded to fill the vacated niche, whereas *Flavobacterium* and *Pseudomonas* remained the dominant taxa in complete SynCom cultures (**Fig. 2b**). *Rhodotorula* abundance, assessed by ITS4 reads, was not significantly altered by the absence of *Pseudomonas* and remained the most prevalent yeast across all dropout combinations. These data indicate that removing *Pseudomonas* primarily affects bacterial rather than yeast composition.

**Fig. 2.**
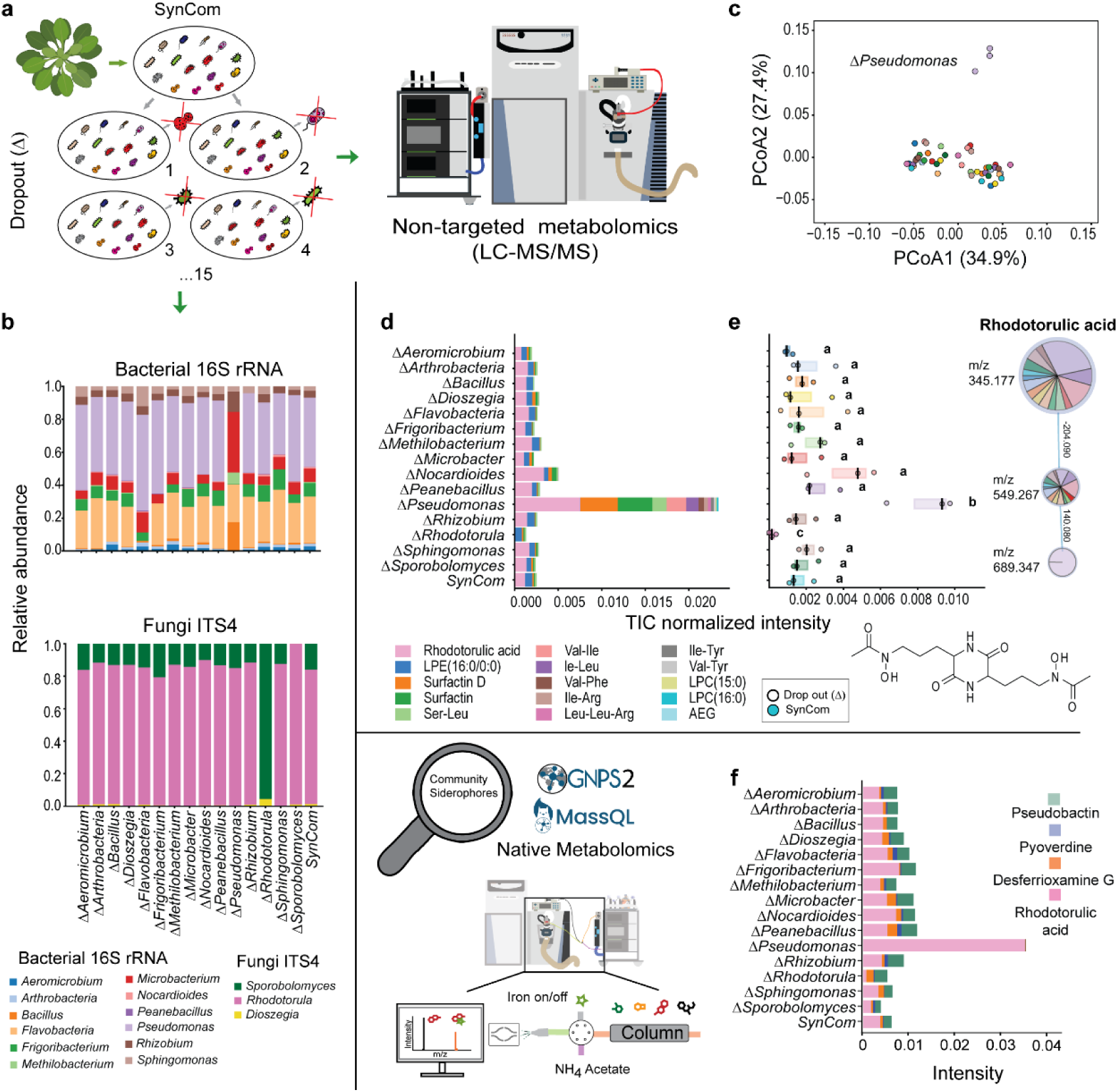
Molecular mechanisms behind the *Rhodotorula*-*Pseudomonas* positive correlation in the “core” microbial community. **a,** Schematic representation of the dropout SynCom co-cultures assembled to investigate the chemical space occupied by each strain within the “core” microbiome. **b,** Relative abundance of strains in dropout SynCom co-cultures, assessed by 16S rRNA/ITS4 Nanopore amplicon sequencing from MM9 agar after 5 days of incubation at 22 °C. **c,** PCoA plot of dropout SynCom metabolomic profiles, samples colored by dropout. **d,** Absolute stacked bars show mean TIC values of the 15 annotated metabolites with the highest ANOVA F-statistics across groups. **e,** Boxplot of TIC-normalized intensity of the annotated node corresponding to rhodotorulic acid, identified from non-targeted metabolomics combined with Feature-Based Molecular Networking (FBMN). Different letters denote significant differences (ANOVA with Tukey’s test, *P* < 0.05). In this network, nodes represent detected features connected by MS² spectral similarity (cosine > 0.75), and edges indicate the corresponding delta-mass relationships. Node size reflects differences in feature intensity across groups. **f,** To focus on metabolites with iron-binding capacity within the community, GNPS molecular networking, MassQL queries, and metal infusion–based validation were applied to identify siderophores and assess their distribution across dropout SynCom extracts. Stacked bar plots display the relative abundance of the main siderophores identified by MassQL and/or metal-infusion native metabolomics for each dropout SynCom co-culture. Peak areas corresponding to nodes within the same cluster and with similar retention times were merged to obtain the most reliable estimates of the different siderophore concentrations among the group.

Across all co-cultures, 5663 LC-MS/MS features were detected after preprocessing, of which 763 differed significantly among dropout combinations (ANOVA, Benjamini/Hochberg False Discovery Rate (FDR) corrected *P* < 0.05; (**data S1)**. Principal coordinate analysis (PCoA) of dropout SynCom metabolomic profiles revealed distinct separation, with Δ*Pseudomonas* clearly clustering from the other groups (**Fig. 2c**). Mean total ion chormatogram (TIC) intensities of the top 15 annotated metabolites with the highest ANOVA F-statistics indicate that RA is the most strongly differentiating metabolite among groups (**Fig. 2d**). RA abundance increased markedly in Δ*Pseudomonas* but was undetectable in *ΔRhodotorula* cultures (*P* = 3.37× 10^⁻7^) (**Fig. 2e**), consistent with the fungal origin of this siderophore and the capacity of gram-negative bacteria such as *Pseudomonas* to scavenge hydroxamate ligands of fungal origin.

Because RA was identified as the metabolite most responsive to *Pseudomonas* removal, we examined siderophore production patterns in greater depth. In addition to compounds annotated via GNPS, we employed MassQL queries and metal-binding native metabolomics to uncover additional iron-chelating molecules (**Extended Data Fig. 2, and Supplementary Table S1**). RA was not detected by MassQL, likely due to its trimeric Fe2RA3 complexation, but was readily confirmed by the metal-infusion approach. When members encoding siderophore biosynthetic gene clusters (BGCs; identified by antiSMASH) were removed, the diversity of detected iron-binding compounds decreased, consistent with intensified iron competition driven by siderophore-producing taxa. Conversely, excluding strains lacking siderophore BGCs increased siderophore diversity, suggesting that, when present, these taxa constrain siderophore producers through niche occupation and competition for iron, thereby limiting the diversification of siderophore profiles (**Extended Data Fig.2)**.

Raw peak-area analysis across all combinations revealed that *Pseudomonas koreensis* produces pseudobactin and an uncharacterized low-weight pyoverdine siderophore, which stimulated other community members to secrete desferrioxamine G and RA (**Fig. 2f**). In the Δ*Pseudomonas* SynCom, RA dominates siderophore production, whereas *Pseudomonas* presence promotes diversification through iron competition. Taken together, these data demonstrate that removing *Pseudomonas* disrupts the community’s metabolic equilibrium, resulting in compensatory production of the fungal siderophore RA. This cross-kingdom metabolic response suggests that *Pseudomonas* not only modulates iron competition but can also exploit siderophores secreted by other SynCom members, providing a mechanistic foundation for cooperative iron acquisition and microbiome stability.

### Rhodotorulic acid selectively stimulates commensal but not pathogenic *Pseudomonas*

Our previous data indicate that the cell-free supernatant from *Rhodotorula kratochvilovae* cultures supports the growth of *Pseudomonas* in isolation (**Extended Data Fig.1**). We first confirmed that the supernatant used in this experiment is enriched in RA (**Extended Data Fig. 3a**). We then tested whether RA can support the growth of the SynCom member *Pseudomonas koreensis*. Supplementation of iron-limited MM9 medium with increasing concentrations of RA significantly stimulated the growth of the *P. koreensis* strain in a dose-dependent manner over 8–9 hours of incubation (**Extended Data Figs. 3A-B**). We also performed growth assays in MM9 supplemented or not with RA in the presence or absence of either glucose or amino acids (aa) to rule out the possibility that *P. koreensis* uses RA as a carbon or nitrogen source (**Extended Data Fig.4**). Altogether, our data indicate that RA enhances bacterial proliferation under iron-limited conditions by providing chelated Fe³⁺. Carbon limitation increases growth rate but simultaneously increases iron demand, making siderophore uptake a limiting factor.

To test the specificity of this interaction, we extended the analysis to additional *Pseudomonas* isolates representing both commensal and opportunistic pathogenic lifestyles. Growth stimulation by *Rhodotorula*-conditioned medium was consistently observed among commensal strains, whereas none of the pathogenic isolates, including those frequently associated with the *Arabidopsis* phyllosphere, showed a significant response compared with untreated controls (**Fig. 3a**). These results demonstrate that RA acts as a selective xenosiderophore, accessible to commensal *Pseudomonas* but unavailable to pathogenic counterparts. By exploiting fungal siderophores, commensal *Pseudomonas* spp. gain an iron-dependent fitness advantage that may restricts opportunistic pathogens.

**Fig. 3.**
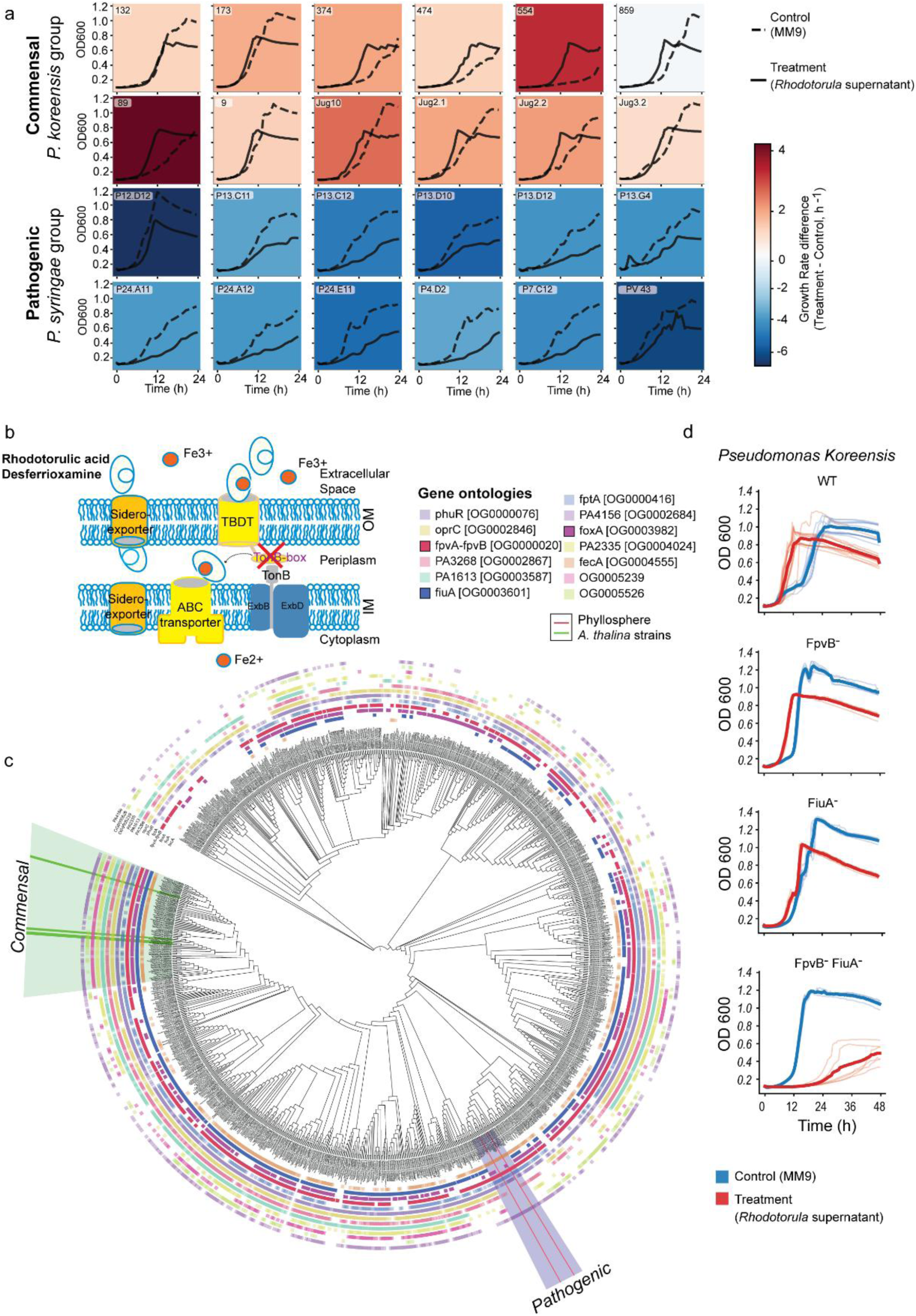
Pangenome analysis reveals absence of specific TonB boxes for phyllosphere core siderophores in pathogenic strains. **a,** Growth curves of 12 commensal and 12 pathogenic *Pseudomonas* strains used in this study. All strains were grown in biological replicates in MM9 medium (dashed lines; control) or MM9 supplemented with 50% (v/v) *Rhodotorula*-conditioned medium (solid lines; treatment). The background heatmap indicates the difference in growth rate between control and treatment conditions, with the color gradient reflecting the magnitude of the effect. **b,** Schematic representation of siderophore transport in Gram-negative bacteria, highlighting the TonB box, which is essential for siderophore translocation into the periplasmic space. **c,** Distribution of TonB-dependent transporters (TBDTs) across the *Pseudomonas* genus, shown in the context of phylogenetic relationships. Annotated TBDTs and those differentially present between commensal (*P. fluorescens* complex) and pathogenic (*P. syringae* group) genomes are indicated. **d,** Deletion of TBDTs in a commensal *Pseudomonas* SynCom member abolishes the growth-promoting activity of the *Rhodotorula* supernatant.

### TonB-dependent transporters mediate RA uptake in commensal *Pseudomonas*

To uncover the genetic basis of RA utilization, we compared the genomes of commensal and pathogenic *Pseudomonas* isolates. Whole-genome sequencing and pangenomic analyses revealed pronounced differences in TonB-dependent transporter (TBDT) repertoires between lifestyle groups (**Extended Data Fig. 5a**). Genes encoding outer-membrane TBDTs with TonB-box signal sequences, which are essential for siderophore translocation, were consistently enriched in commensal strains but absent from pathogenic ones (**Fig. 3b**). These included gene clusters homologous to *fiuA,* which mediate the uptake of hydroxamate siderophores such as RA, as well as *foxA* involved in the transport of desferrioxamine (**Extended Data Fig. 5B**).

To determine whether these patterns extend across the genus *Pseudomonas*, we expanded our analysis to include all available species, revealing a consistent absence of these TBDTs within the *P. syringae* clade (**Fig. 3c**). Focusing further on dominant *Pseudomonas* species associated with wild Arabidopsis, we screened ∼1400 publicly available *Pseudomonas viridiflava* genomes for Pfam-annotated domains required for functional siderophore uptake. This analysis showed that the genes encoding the TonB-box-containing transporters *fiuA* and *foxA* were consistently absent from pathogenic *P. viridiflava* genomes, whereas all three TBDTs were retained in leaf-isolated commensal strains (**Extended Data Fig. 6**).

To assess the role of TBDTs in siderophore uptake, we generated knockout derivatives of the SynCom commensal *P. koreensis* and monitored their growth in the presence of *Rhodotorula* supernatant ^16^. Deletion of *fpvB^-^*, also involved in RA uptake, had no significant effect, whereas deletion of *fiuA^-^* reduced growth promotion. The double mutant lacking both *fiuA^-^* and *fpvB^-^ (fiuA^-^fpvB^-^)* failed to benefit from *Rhodotorula* supernatant, confirming that both receptors are essential to sustain the growth advantage observed in the wild type (**Fig. 3d**). The growth rate *fiuA^-^fpvB^-^* was comparable to that of the pathogenic *P. viridiflava* strains (**Fig. 3a**). Importantly, iron supplementation rescued the growth defect of the double mutant in the *Rhodotorula* supernatant (**Extended Data Fig.7).** These data suggest iron competition and capture is mediated through TBDTs, rather than carbon availability, as the mechanism underlying the differential responses of commensal and pathogenic *Pseudomonas* strains to *Rhodotorula* conditioned medium.

### Rhodotorulic Acid indirectly shapes the microbiome through plant iron stress responses

Because RA accumulated when commensal *Pseudomonas* were absent or lacked the corresponding transport system, we next examined how *Pseudomonas*, *Rhodotorula*, the siderophore RA, and their combinations influence the metabolome of sterile plants following infection by the pathogen *P. syringae* (**Extended Data Fig. 8A**). We speculated that interactions between *Rhodotorula* and *Pseudomonas* may confer indirect host protection by creating an iron-limiting environment that excludes pathogenic microbes. Non-targeted metabolomics of plant aerial tissue revealed that concomitant exposure to *Pseudomonas* and *Rhodotorula* or to Pseudomonas and RA interactions was associated with depletion of jasmonic acid (JA) precursors **(Extended Data Fig. 8**) relative to control and *Pseudomonas* only treatments. JA plays a central role in plant defense and is tightly linked to lignin biosynthesis, a pathway critical for cell-wall reinforcement and stress tolerance ^17^. Consistently, multiple lignin precursors were upregulated ^18^, suggesting a coordinated host response to iron deficiency that enhances structural defenses while facilitating iron mobilization ^19^. Among these compounds, sinapic acid, 3,5-dimethoxy-4-hydroxycinnamic acid, and sinapoylmalate were significantly upregulated under both RA and *Rhodotorula* treatments (**Extended Data Figs. 8a and 9**). Notably, sinapic acid was further detected to be metabolized by several SynCom core microbiome members, including *Pseudomonas, Sphingomonas, Nocardiodes, Dioszegia, and Bacillus,* suggesting a potential metabolic link between host defense metabolites and microbial community dynamics (**Extended Data Fig. 8b**).

### Role of *FiuA* and *FpvB* in plant protection against *Pseudomonas syringae*

To investigate whether TBDTs in *P. koreensis* underlie *Rhodotorula*-mediated plant protection in the context of the SynCom, we generated community derivatives in which the wild-type commensal *Pseudomonas* was replaced by each of the TBDT knockout mutants. These modified SynComs were inoculated onto gnotobiotic *A. thaliana* to assess loss or reduction of the protective phenotype compared to wild type (WT) SynCom. Following a 5-day establishment period, plants were infected with *P. syringae,* disease progression was quantified using both the percentage of symptomatic leaves per plant and Colony-forming units (CFUs) per gram of fresh plant weight (CFU g⁻¹ FW). A clear gradient in protection emerged plants inoculated with the *fpvB* knockout SynCom exhibited disease levels comparable to the wild type community, whereas, the SynCom harboring Pseudomonas *fiuA-fpvB-* showed the highest infection levels, indistinguishable from the non-inoculated control (**Fig. 4a**).

**Fig. 4.**
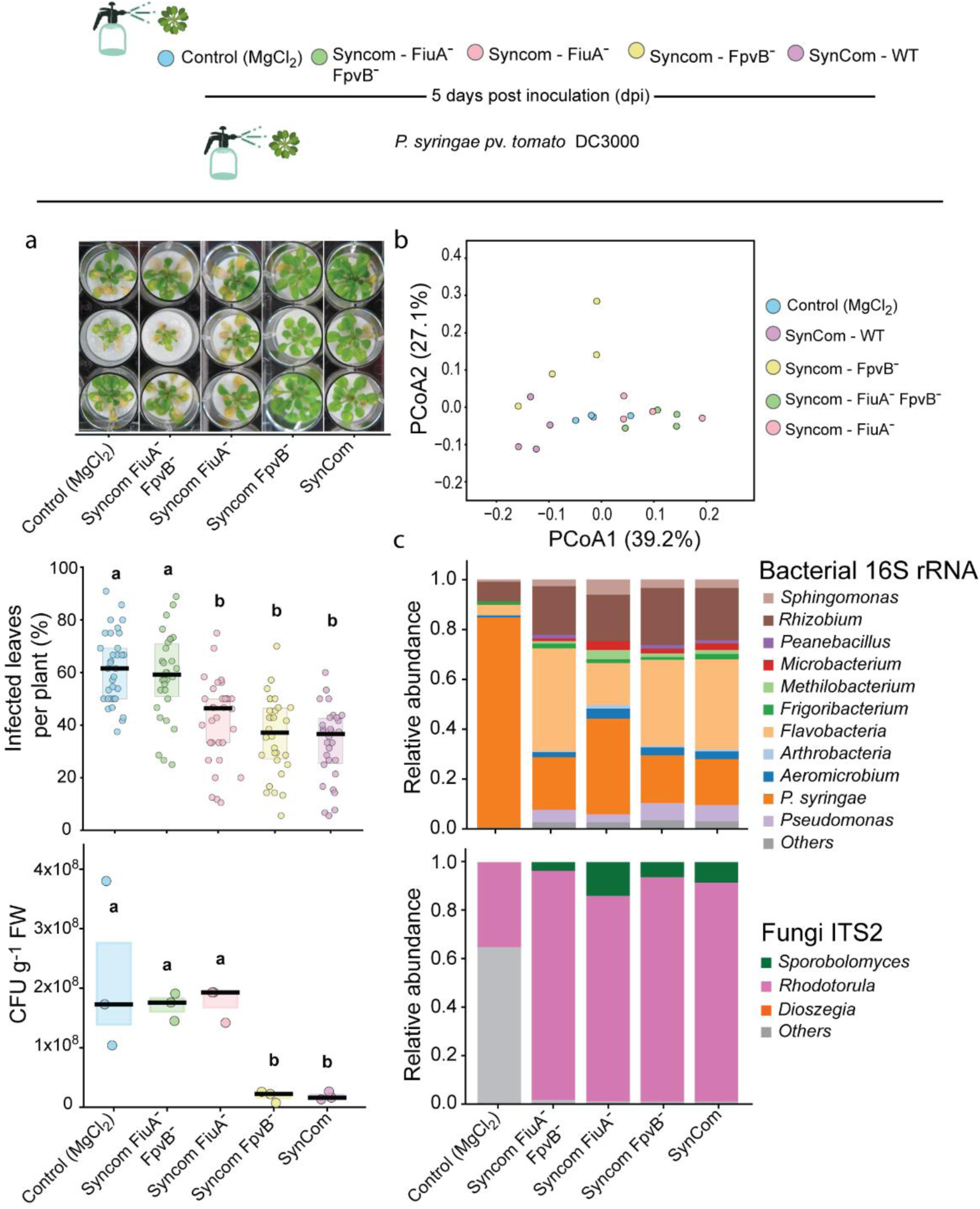
Interkingdom interactions promote plant protection. *A. thaliana* plants were grown sterile condition and treated with 10 mM MgCl₂ containing either the complete SynCom or derivative communities in which *P. koreensis* was replaced by knockout strains generated in this study. Five days later, all plants, including controls, were inoculated with *P. syringae* at equal optical density. **a,** Disease severity, quantified as percentage of infected tissue and pathogen load (CFU g⁻¹ fresh weight). Different letters denote significant differences (ANOVA with Tukey’s test, *P* < 0.05). **b,** Principal coordinates analysis (PCoA) of non-targeted metabolomics data, based on Bray–Curtis distances of TIC-normalized features. Points represent individual samples; colors indicate treatments. Metabolite profiles differed significantly among SynComs (PERMANOVA, pseudo-F = 0.01, *P* = 0.026). **c,** Relative abundances of SynCom members and P. syringae from epiphytic microbial DNA, including bacterial and fungal fractions.

Non-targeted metabolomics of plant aerial tissues corroborated these patterns. Principal coordinates analysis (PCoA) and hierarchical clustering revealed that plants colonized with the *fpvB* knockout SynCom displayed metabolomic profiles similar to those receiving the wild-type community, while plants inoculated with *fiuA* and the double mutant SynCom clustered with the control samples (**Fig. 4a and fig. S10**). Our results show that polyunsaturated fatty acids and oxylipids (peroxidized fatty acids), direct precursors of jasmonic acid, are differentially regulated across treatments, in line with previous reports implicating jasmonic acid in iron-homeostasis responses (**Extended Data Figs. 8a-b**). Lignin precursors levels increased under conditions where iron availability may have been limited, potentially as a consequence of increased microbial competition for iron. Amplicon sequencing of the epiphytic microbial community confirmed that all *Pseudomonas* mutant strains persisted stably within the SynCom following plant inoculation (**Fig. 4c**). In control plants, which had not been exposed to any microbial community, a small number of reads were detected, likely reflecting minor cross-contamination. Together, these results demonstrate that TBDTs link microbial iron acquisition to host metabolic response, thereby shaping community-level protection against *P. syringae*.

## Conclusion

Iron is a limiting micronutrient, and its reduced availability restricts pathogen colonization in many microbiomes, including the phyllosphere. Keystone microbial species, such as commensal *Pseudomonas* members, stabilize the phyllosphere. Disruption of plant-associated microbiomes compromises plant productivity, ecological stability, and stress resistance, underscoring the importance of maintaining microbiome integrity. Yet, the mechanisms underlying the protective effects of *Pseudomonas* mediated by microbial interactions remain largely unresolved. Using a synthetic community derived from natural *A. thaliana* microbiota and using state-of-the-art analytical tools, we show that *Rhodotorula* enhances the growth of commensal *Pseudomonas* through the uptake and utilization of RA, a process dependent on the exclusive presence of a TonB-dependent transporter (TBDT). Together with other community siderophores, such as desferrioxamine G, this interaction reduces iron availability in the environment, thereby shaping microbial interactions and influencing host-microbiome dynamics. In contrast, members of the pathogenic *P. syringae* group are unable to utilize xenosiderophores in the community, as confirmed by pangenome and *in-vitro* analysis (**Fig. 3b**). Commensal *Pseudomonas* produce their own siderophores, thereby intensifying competition for iron and limiting its availability within the plant microbiome. This severe iron scarcity reduces the survival capacity of pathogenic microbes, directly enhancing host protection. Furthermore, we found that spraying apo-rhodotorulic acid (apo-RA) triggers a host stress response, leading to the activation of the lignin biosynthesis pathway, which our results suggest is beneficially exploited by members of the core microbiome (**Extended Data Fig. 8b**). As such, our study identifies iron-mediated microbial competition as a key process in protecting and stabilizing plant microbiomes (**Fig. 5**).

**Fig. 5:**
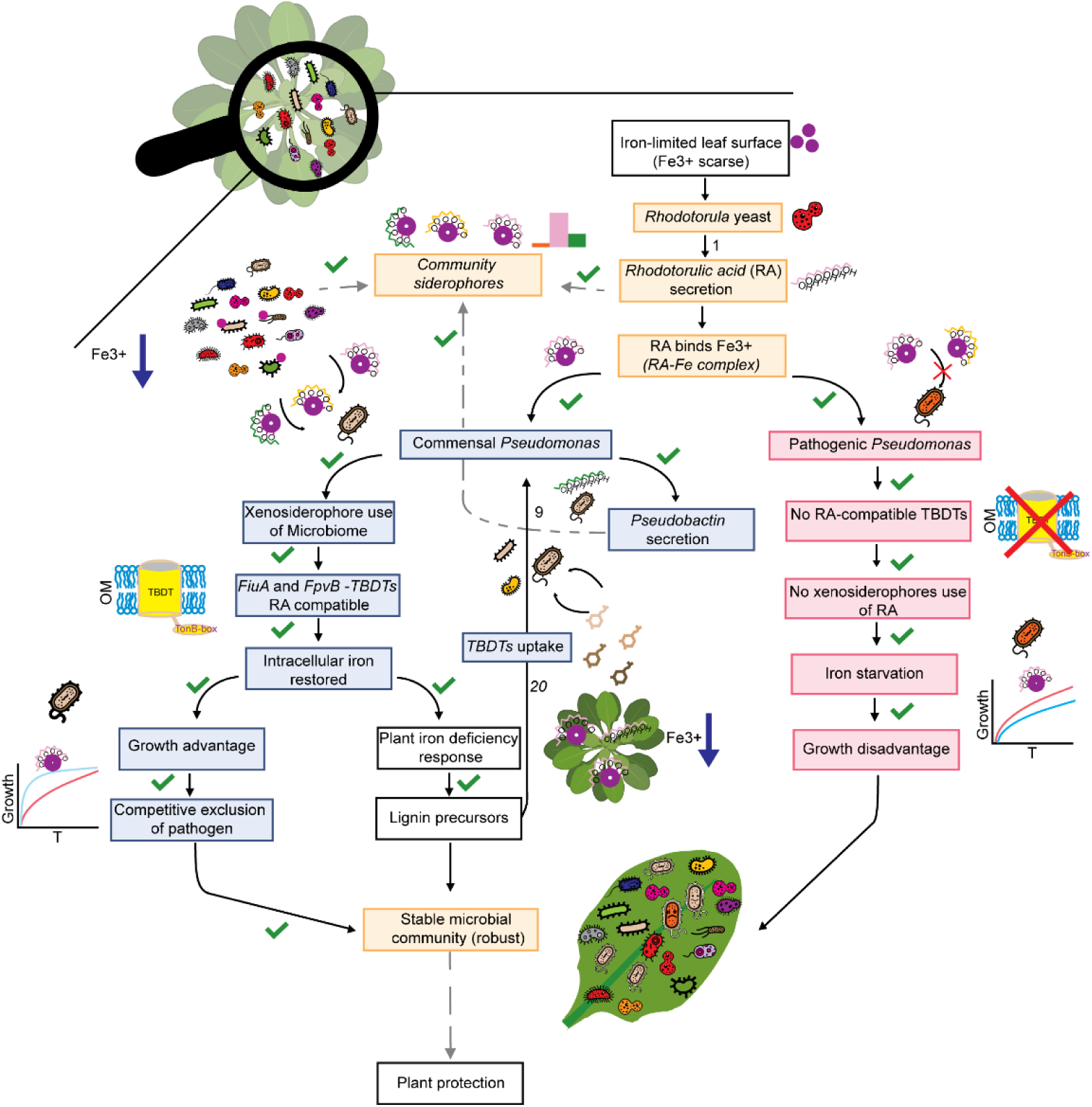
Siderophore-mediated cooperation in phyllosphere iron acquisition promotes plant protection. Flowchart depicting the major mechanisms identified in this study. ^9,20^

Our findings extend the current understanding of interkingdom interactions in plant-associated microbiomes by demonstrating their role in enhancing host protection within the phyllosphere. While previous studies have documented beneficial interkingdom interactions in the rhizosphere ^21^, similar dynamics in the phyllosphere remain underexplored. This is particularly relevant given the established role of the phyllosphere as a critical front line in plant defence against foliar pathogens ^22^. Commensal *Pseudomonas* strains, such as those included in our synthetic community, suppress the growth of opportunistic *Pseudomonas* pathogens that are widely distributed across agricultural and wild *A. thaliana* populations ^10^. *M*embers of the genus *Pseudomonas* can act as both beneficial and pathogenic members of the phyllosphere microbiota ^23^. Iron acquisition has emerged as a key mechanism by which commensal *Pseudomonas* limit pathogen proliferation ^11^. In our system, the presence of yeast-derived siderophores, specifically RA produced by *Rhodotorula*, promotes directly commensal *Pseudomonas* growth while excluding pathogenic strains. We observed that the simple siderophore triggers plant immunity and lignin biosynthesis, with a beneficial effect on members of the core microbiome, thereby indirectly stabilizing the community. These selective effects may facilitate niche dominance, reducing pathogen access to essential resources ^24–26^. Pathogen success depends not only on host susceptibility and nutrient availability, but also on the capacity to compete within a complex microbiome ^27–29^.

Our work shows that interkingdom cooperation in iron utilization based on the presence of TBDTs in keystone members, such as commensal *Pseudomonas* in the phyllosphere, shapes both plant health and microbial community structure, revealing an underappreciated layer of plant defence. By promoting beneficial microbes and enhancing stress resilience, this mechanism may help stabilize plant populations under anthropogenic pressures. Harnessing such interactions offers a promising path toward microbiome-informed strategies for sustainable agriculture and food system resilience.

## Methods

### Rhodotorula Correlation Network

Co-abundance network analysis was performed to investigate the interactions of *Rhodotorula* with other microbes in the phyllosphere microbiome of *Arabidopsis*. Operational Taxonomic Units (OTUs) were directly obtained from Mahmoudi et al.(2024) ^30^. The OTU table for all samples (epiphytes and endophytes), including bacterial and eukaryotic (fungal and nonfungal) OTUs, was filtered to retain only those OTUs present in at least five samples with more than ten reads. SparCC ^31^ correlations were computed using the FastSpar ^32^ platform with default parameters. Permuted *p*-values for each correlation were derived from 1,000 bootstrap datasets, and only correlations with *P* ≤ 0.05 were retained for further analysis. The co-abundance network was visualized using Cytoscape (version 3.10.0) ^33^. Nodes (OTUs) in the network were labelled at the genus level based on OTU assignments using the UNITE ^34^ database (version 02_02_2019) for fungi and the SILVA ^35^ database(version 138.1) for bacteria, as described in Mahmoudi et al. 2024 ^30^.

### Dropout co-cultivation of the “Core” SynCom Members

In this study, the SynCom, comprised of a diverse array of bacteria and yeast strains ^36^, was meticulously cultivated in co-culture assembled by dropping out of the members besides a full Syncom used as a co-culture positive control. Initially, strains were revived from 30% glycerol (≥ 99.5%) cryo-stocks and cultivated on specific agar plates - Nutrient Broth (NB) agar for bacteria and Potato Dextrose Agar (PDA) agar for yeast. After incubating at 22 °C for 3 days, the strains were transitioned into liquid cultures, utilizing NB and Potato Dextrose Broth (PDB) for bacteria and yeast, respectively, maintaining the same incubation conditions for an additional 2-3 days.

Following this, the bacteria and yeast liquid cultures were centrifuged at 4000 x g, and the resulting pellets were washed thrice with an MgCl_2_ 10 mM solution. Subsequently, the strains were suspended in MgCl_2_ 10 mM, yielding solutions with an optical density (OD) absorbance 1 OD. These solutions were then utilized to generate 16 distinct co-cultures, each excluding one of the 15 SynCom strains, a SynCom containing all members for comparative analysis was also included in these co-cultures. These co-cultures were inoculated onto minimal medium (MM9) agar plates and allowed to grow at 22 °C for 5 days. After that, microbial cultures were harvested by carefully scraping cells from the plates.

### Amplicon sequencing of in-vitro SynCom communities with dropout members

DNA was extracted using the ZymoBIOMICS DNA Miniprep Kit (Zymo Research Corp., Irvine, CA, USA) following the manufacturer’s protocol. For bacterial community characterization, 16S amplicons of the V3-V8 region (∼1,300 bp) were generated by PCR using primers 341F ^37^ and 1378R ^38^ based on the protocol previously described by Seol et al ^39^. Reactions for generating 16S amplicons were carried out with 50 ng of DNA in 11.5 μL of nuclease-free water, 0.5 μL of each primer and 12.5 μL of LongAMP Taq 2x Master Mix (New England Biolabs, Ipwich, MA, USA) for a total reaction volume of 25 μL. The PCR cycle program was: 1 minute 95 °C; 25 cycles of 20 s 95 °C, 30 seconds at 55 °C, 120 seconds at 65 °C and a final elongation time of 5 minutes at the same temperature. For fungi characterization, ITS amplicons were generated using primers ITS1 TCCGTAGGTGAACCTGCGG and ITS4 TCCTCCGCTTATTGATATGC ^40^. Reactions for generating ITS amplicons were carried out with 50 ng of DNA in 11.5 μL of nuclease-free water, 0.5 μL of each primer and 12.5 μL of LongAMP Taq 2x Master Mix (New England Biolabs, Ipwich, MA, USA) for a total reaction volume of 25 μL. The PCR cycle program was: 1 minute 95 ° C; 25 cycles of 15 seconds 95 °C, 15 seconds at 55 °C, 30 seconds at 65 °C and a final elongation time of 2 minutes at the same temperature. After PCR, Amplicons were purified using AMPure XP beads (Beckman Coulter, Brea CA, USA) at a 0.6 x bead-to-sample ratio and quantified with a Qubit 4 fluorometer. Nanopore sequencing libraries were prepared with the Rapid Barcoding Kit (SQK-RBK114.96) using the Amplicon sequencing from DNA protocol (available at https://nanoporetech.com/document/rapid-sequencing-v14-amplicon-sequencing-sqk-rbk114-24-or-sqk) and sequenced on MinION V14 flow cells (Oxford Nanopore Technologies) using MinKNOW v25.05.14. Only reads with a quality score ≥10 were retained and demultiplexed. Taxonomic classification was performed using Emu v3.4.5 ^41^ with a custom rRNA/ITS operon reference database generated from the species genomes using the in silico PCR implemented in EMBOSS-primersearch ^42^ as previously reported ^39^.

### LC-MS/MS non-targeted metabolomic analysis

The extraction of metabolites involved the addition of an 80% methanol solution (LC/MS grade, Fisher Scientific) to the microbial cultures. The solutions were then vortexed for 2 minutes followed by ultrasonication for 15 minutes in an ultrasonicator bath. Subsequently, debris was removed from the solution through centrifugation at 13000 x g for 20 minutes.

Quantification of metabolites was achieved by evaporating the solution in pre-weighted vials. To create a 5 mg/mL concentration, the dried metabolites were dissolved in a 50% methanol solution. Non-target metabolomics was performed by liquid chromatography-tandem mass spectrometry (LC-HRMS/MS) with a UHPLC coupled to a Q-Exactive HF mass spectrometer. In short, the UHPLC separation was performed using a C18 core-shell column (Kinetex, 50 × 2.1 mm, 1.7 µm particle size, 100 A pore size, Phenomenex, Torrance, USA). The mobile phases used were solvent **a,** H_2_O (LC/MS grade, Fisher Scientific) + 0.1% formic acid (FA) and solvent **b,** acetonitrile (LC/MS grade, Fisher Scientific) + 0.1% FA. After the sample injection, a linear gradient method of 10 min was used to elute small molecules, and the flow rate was set to 500 µl/minute (normal flow). The following separation conditions were set: in the time range 0 - 8 minutes from 5% to 50% solvent **b,** was used, 8 – 10 minutes from 50 to 99% B, followed by 3 minute washout phase at 99% B and 3 minute re-equilibration phase at 5% B.

The measurements were conducted in positive mode, the Heated Electrospray Ionization (HESI) parameters included a sheath gas flow rate of 50 L/minute, auxiliary gas flow rate of 12 L/minute, and sweep gas flow rate of 1 L/minute. The spray voltage was set to 3.50 kV and the inlet capillary temperature to 250 °C, while the S-lens RF level to 50 V and the auxiliary gas heater temperature to 400 °C. The full MS survey scan acquisition range was set to 120–1,800 m/z with a resolution of 45,000, automatic gain control (AGC) of 1E6, and maximum injection time of 100ms with one micro-scan. DDA MS/MS spectra acquisition was performed in data-dependent acquisition (DDA) mode with TopN set to (5) as a consequence the five most abundant precursor ions of the survey MS scan were destined for MS/MS fragmentation. The resolution of the MS/MS spectra was set to 15,000, the AGC target to 5E5, and the maximum injection time to 50 ms. The isolation window for the precursor ion selection was set to 1.0 m/z and the normalized collision energy was set to a stepped increase from 25 to 35 to 45. MS/MS experiments were triggered at the apex of peaks within 2–15 seconds from their first occurrence. Dynamic exclusion was set to 5 seconds ^43^.

Raw data were converted into centroid mode file mzML, and uploaded on MZmine ^44^ for spectra processing. Briefly, the mass levels of noise were set at 3.0E5 and 1.0E4 for the mass detection at MS1 and MS2, respectively. The ADAP Chromatogram Builder Module was used with at least 5 consecutive scans needed for the peak definition and maximum mass tolerance set at 10 ppm. To achieve the feature resolving the ms/ms scan pairing and the retention time dimension were activated. In addition, the isotopic peak finder and the join aligner modules were applied together with the export modules for FBMN (batch file in the SI). Feature table and mgf file were used to create FBMN on the GNPS2 platform ^45^. FBMN task ID was used for data uploading on FBMN-STAT for data preparation, including blank removal, imputation, and normalization ^46^. Statistical analyses were conducted using custom Python code in Jupyter Notebook.

### Native metabolomics with the metal infusion

To investigate the dynamics of siderophore production within the SynCom, we followed the method established by Aron et al. (2022) ^47^. Extracts from the dropout co-culture experiments, conducted in biological triplicate, were analysed to detect the presence of iron-binding metabolites, across all 15 dropout co-cultures. The analysis employed a two-step native ESI-LC-MS/MS workflow, which involved post-column infusion of ammonium acetate buffer at an initial concentration of 100 mM for post-pH adjustment, followed by the infusion of Iron(III) chloride hexahydrate solution at an initial concentration of 3 mM. Data acquisition was performed in data-dependent acquisition (DDA) mode, adhering to the parameters outlined in the LC-MS/MS non-targeted metabolomic analysis section described previously. Data analysis was conducted using the ion identity molecular networking (IIMN) workflow to observe the iron-bound adduct, in combination with MZmine4 and the GNPS platform. The converted raw data files coming from the metal infusion “Off” samples in centroid mode (mzML format) were additionally utilized for in silico siderophore detection. MassQL queries were formulated and used in the workflow called “massql_workflow” available on the GNPS2 platform ^48^. This allows us to identify iron-bound small molecules within our dataset. The queries considered both single-charged and double-charged iron masses. Cytoscape (version 3.7.1)^33^ was used to visualize molecular networks.

### Leaf-associated Bacteria Isolation and Characterization

Commensal *Pseudomonas* strains from the *Arabidopsis* leaf–associated culture collection (Kemen laboratory), stored in 30% glycerol (≥ 99.5%) at −80 °C, were used in this study. These strains were previously subcultured and confirmed for purity. The 16S rRNA gene was PCR-amplified, sequenced, and taxonomically assigned via BLAST against the NCBI Type Strain database. Pathogenic *Pseudomonas* strains were provided by the Weigel laboratory (Max Planck Institute for Biology, Tübingen), and their genome sequences are publicly available.

### Whole Genome Sequencing (WGS) and taxonomy of Pseudomonas strains

Commensal *Pseudomonas* leaf-associated bacteria were cultured in NB broth and genomic DNA from each was isolated, and the quality of the genomic DNA was assessed using agarose gel electrophoresis. DNA concentrations were estimated using a Nanodrop spectrophotometer ND-100 (Thermo Fisher Scientific, USA) and Quantus™ Fluorometer (Promega). Both the ratio A260/280 and gel electrophoresis were used to ascertain the quality and purity of DNA samples. The input of 1 μg of genomic DNA was used for library construction and sequencing of libraries Illumina platform (Illumina, San Diego, CA) using a NovaSeq X Plus, 2 x 151 bp* paired-end kit in the sequencing facility in CeGaT (Tübingen) (**Supplementary Table 2**).

Raw sequencing reads of *Pseudomonas* strains were first demultiplexed and subjected to quality control and adapter trimming using fastp (v 0.23.4) ^49^with default parameters. The trimmed reads were then assembled de novo using SPAdes (v 4.0.0) ^50^. Genome annotation was performed using Bakta (v 1.9.4) ^51^ with database version 5.1. Taxonomic classification of the assembled genomes was conducted using GTDB-Tk (v 2.6.1) ^52^ against GTDB database release r226. Functional annotation of identified CDSs was done using eggNOG-mapper (v 2.1.12) ^53^ in DIAMOND mode.

### Pangenome analysis

To analyse the pangenome of the assembled genomes of the isolates, as well as the reference genomes of *P. atacamensis* Jug 3.2 (GCF_022802935.1)*, P. koreensis* (GCF_001654515.1)*, P. syringae* (GCF_018394375.1), and *P. viridiflava* (GCF_900184295.1) in anvi’o, raw genome assemblies were first reformatted keeping contigs of at least 2,500 bp. Several annotation steps were performed, including the assignment of functional annotations with NCBI COGs (anvi-run-ncbi-cogs), KEGG ortholog annotations (anvi-run-kegg-kofams), PFAM domain identification (anvi-run-pfams). The pangenome analysis was refined using an MCL inflation value of 10 and a minimum bit score ratio of 0.5, with NCBI BLAST as the sequence similarity search tool. Additional metadata regarding the pathogenicity of the isolates was added to the anvi’o pangenome analysis, with *P. viridiflava* and *P. syringae* strains marked as positive for pathogenicity and the rest marked negative. Finally, functional enrichment analyses were conducted using anvi-compute-functional-enrichment-in-pan, focusing on genes associated with pathogenicity based on COG20_FUNCTION and COG20_PATHWAY annotations, and the results were visualized using anvi’o in interactive mode.

The gene-enrichment analysis of the pathogenic-vs-commensal strains revealed different clustering of COG4773 (FhuE) and COG4774 (Fiu) functions. To investigate these further, we build a more comprehensive pangenome of all representative species available in the GTDB database, using anvi’o as described above. The presence-absence of gene clusters associated with these functions were then visualized using iTOL, projected onto the subtree of the genus *Pseudomonas_E* from GTDB.

In order to further investigate the functional diversity of TonB-related protein in *P. viridiflava* strains, we screened for proteins containing the PFAM domains PF00593.30 (TonB dependent receptor-like, beta-barrel), PF07660.19 (Secretin and TonB N terminus short domain (STN)), or PF07715.20 (TonB-dependent Receptor Plug Domain) in all publicly available *P. viridiflava* genome assemblies (1396 genomes from GTDB). We collected the respective HMM models from PFAM and searched them against *P. viridiflava* protein sequences using hmmscan and extracted all the proteins that hit a combination of these domains (6051 proteins). We generated a multiple sequence alignment of these proteins using MAFFT (v 7.526, with parameters --localpair --maxiterate 1000) ^54^, and followed by a maximum likelihood tree construction using FastTree (v 2.1.11, with parameters --gamma) ^55^.

### Construction of Pseudomonas koreensis knockout strains

Chromosomal deletions were performed by single homologous recombination using two plasmids with different antibiotic cassettes. Briefly, 530 bp of fpvB (RS21125), and 597 bp of fiuA (RS16190) were cloned into pCR 2.1 and, pJQ200KS plasmids, respectively (Invitrogen). Cloning into pCR 2.1 was performed according to manufacturer recommendations while restriction enzymes and T4-DNA ligase were used for the construct (BamHI-XhoI for pJQ200KS-fiuA). The oligonucleotides used are shown in below. The resulting plasmids were confirmed by PCR, restriction pattern and sequencing, and then electroporated into *Pseudomonas koreensis*. Selection was performed using the corresponding antibiotic resistances (Km, and Gm respectively). Resistant clones were checked by PCR to confirm mutation in the target genes. Once single mutants were obtained, we next performed double mutants by cloning plasmids into single mutant strains following the same procedure described above.

Oligos:

FiuA_PJQ_fwd GGATCCCACCAAACTCACTCTGCTGAGCCAGTTCAC

FiuA_PJQ_rev CTCGAGGTCGAGAGCCATTTGGTCCTGCACGTAGAG

Fpvb_pCR2.1_fwd TCACGGCCTATTCTACGCCATCGGTGAAGC

Fbvb_pCR2.1_rev GACTTGCCGCTACGCACGAAGTCAGGCTTG

### SynCom and Pseudomonas growth curves under Rhodotorula supernatant

The 15 SynCom member strains were used to generate individual growth curves in the presence of *Rhodotorula* culture supernatant. Treatment samples consisted of a 1:1 (v/v) mixture of supernatant from a *Rhodotorula* strain cultured for 5 days in MM9 medium and fresh MM9 medium; fresh MM9 medium alone was used as the control. MM9 minimal medium was prepared as described previously ^36^. After autoclaving the basal salts MgSO₄ and CaCl₂, glucose, a defined amino acid mixture (17 essential amino acids), and a trace element solution were added under sterile conditions after autoclaving. Experiments were performed in 96-well plates using a Tecan plate reader. Optical density at 600 nm (OD₆₀₀) was measured every 30 minutes for 24–48 hours at 22 °C with continuous shaking at 125 rpm.

Rhodotorulic acid, a siderophore present in the *Rhodotorula* supernatant, was detected and quantified by using LC/MS-MS analysis. The *Rhodotorula* exometabolome was isolated by solid-phase extraction (SPE), and 400 µL of the crude extract, concentrated to 5 mg/mL, was subjected to fractionation. A rhodotorulic acid standard was obtained from Biophore Research Products (Rottenburg, Germany) and was used as a positive control to assess its effect on bacterial growth through growth-curve experiments. Biological replicates were included for all experiments.

The contribution of rhodotorulic acid to the growth promotion of SynCom *Pseudomonas* strains was evaluated, and additional growth-curve assays were performed using multiple plant-associated *Pseudomonas* strains in the presence of *Rhodotorula* supernatant. To check if *Pseudomonas* grows in varying nutrient conditions and rhodotorulic acid supplementation, growth curves were performed in MM9 medium, which served as the control condition, and seven additional treatments were tested by omitting glucose and/or the amino acid mixture and by supplementing the media with rhodotorulic acid (50 μg/mL). All media were prepared fresh and used for growth curve measurements under identical experimental conditions.

### In-planta protection evaluation assays

For the greenhouse experiments, seeds of *Arabidopsis thaliana* (genotype WS-0) were sown and allowed to germinate in soil under short-day conditions (10 hours of light at 21 °C and 14 hours of darkness at 16 °C). After one month, for non-sterilele plants, the resulting seedlings were individually transplanted into 6-pot trays and cultivated under short-day conditions. Adult plants were divided into experimental groups and treated with the SynCom strains *Pseudomonas*, *Rhodotorula*, or a combination of both. Each treatment was applied by spraying the leaves with a microbial suspension prepared in 10 mM MgCl₂. Control plants were sprayed with 10 mM MgCl₂ solution without microbial inoculation. After five days of microbial establishment on the leaves, both treated and control plant groups were inoculated with *Pseudomonas syringae* pv. *tomato* DC3000. Disease progression was assessed five days after pathogen inoculation by counting the number of symptomatic leaves relative to the total number of leaves per plant.

Gnotobiotic plant experiments were performed to investigate the effects of *Pseudomonas*, *Rhodotorula*, the siderophore rhodotorulic acid (RA), and their combinations on the host, as well as to assess SynComs and SynCom variants containing *Pseudomonas* knockout strains. Experiments were conducted using *Arabidopsis thaliana* WS-0 genotype seeds, which were surface-sterilized prior to use. Seeds were germinated on plant agar plates under short-day conditions (10 hours light at 21 °C, 14 hours dark at 16 °C). Seedlings of uniform size were subsequently transferred individually into 12-well plates for further growth and treatment. Sterile split dishes were used to physically separate the aerial parts of the plants from the agar medium containing the root system. The treated groups received a foliar spray containing a suspension of the 15 strains comprising the SynCom. In the respective treatment groups, the *Pseudomonas* member (wild type) was replaced with the knockout variants (*fpvB⁻*, *fiuA⁻*, or *fpvB⁻ fiuA⁻*). Each strain was adjusted to an OD₆₀₀ of 0.2 in 10 mM MgCl₂ before application. Control plants were sprayed with 10 mM MgCl₂ solution alone. Pathogen inoculation with *P. syringae* pv. *tomato* DC3000 for all groups was performed five days after the SynCom treatment to allow for microbial establishment on the phyllosphere. Plates containing plants were maintained under short-day conditions. Gnotobiotic plants were then assessed for disease symptoms as described for the greenhouse experiments, by quantifying the percentage of symptomatic leaves over the total. In addition, *P. syringae* quantification was determined by counting colony-forming units (CFUs) per gram of fresh weight (FW). The epiphytic microbial community was collected for amplicon sequencing, and additional plants were harvested for metabolite extraction and subsequent high-resolution tandem mass spectrometry (LC-MS/MS) analysis.

### Amplicon sequencing of the epiphytic synCom community

*Arabidopsis thaliana* plants were grown gnotobiotically and inoculated with either the complete SynCom or derivative communities in which *Pseudomonas koreensis* 3.2 was replaced by knockout strains generated in this study, all suspended in 10 mM MgCl₂. Plants were harvested and briefly washed by shaking in 5 mL Milli-Q water for 10 seconds, after which the wash solution was discarded. Subsequently, 5 mL of epiphyte wash solution (1 × TE + 0.1% Triton X-100) was added, and the plants were gently shaken for 1 minute. Epiphytic microorganisms were collected by transferring the resulting suspension into syringe barrels mounted in filter holders. After reinserting the plungers, the suspension was filtered through 0.22 µm mixed cellulose ester (MCE) membrane filters. The filter holders were then opened, and the filters were carefully retrieved and stored in 2mL screw-cap tubes. The retrieved membrane filters were frozen in liquid nitrogen and mechanically disrupted by cryogenic lysis. Samples were processed at 6300 rpm for four 30 second cycles with 15 second pauses between cycles, yielding a finely homogenized material for subsequent extraction steps.

DNA was extracted using the same approach reported above. Genomic DNA was quantified with a Qubit dsDNA BR/HS Assay Kit (Thermo Fisher) and normalized to 5-25 ng Input for library preparation. Library preparation was performed as a two-step PCR approach. The first step PCR for the 16S rRNA V4 region was performed in 15 µl reactions including KAPA HiFi HotStart ReadyMix (Roche), 515F and 806R (Parada et al 2016; April et al 2015) and template DNA (PCR program: 95 °C for 3 minutes, 30x (98 °C for 20 seconds, 55 °C for 15 seconds, 72 °C for 15 seconds), 72 °C for 5 minutes). While the first step PCR for the ITS1 region was performed in 15 µl reactions including KAPA HiFi HotStart ReadyMix (Roche), ITS1F and ITS2R (Gardes and Bruns 1993; White et al. 1990) and template DNA (PCR program: 95 °C for 3 minutes, 35x (98 °C for 20 seconds, 50 °C for 1 minute, 72 °C for 90 seconds), 72 °C for 5 minutes).

PCR products were purified using 12 µl AMPure XP beads and eluted in 26 µL 10 mM Tris-HCl. Indexing for both target regions was performed in the second step PCR including KAPA HiFi HotStart ReadyMix (Roche), index primer mix (IDT® for Illumina Nextera DNA Unique Dual Indexes), purified first PCR product as template (PCR program: 95°C for 3 min, 8x (95°C for 30 sec, 55°C for 30 sec, 72°C for 30 sec), 72°C for 5 min). After another bead purification (14 µl AMPure XP beads, eluted in 15 µL 10mM Tris-HCl) the libraries were checked for correct fragment length on an agarose gel, quantified with a Qubit dsDNA BR Assay Kit (Thermo Fisher) and pooled equimolarly. The pool was sequenced on an Illumina Miseq with a v3 sequencing kit and 2×300 bp read length and a depth of 40-140k reads per sample.

Raw sequencing reads were demultiplexed and quality-trimmed using QIIME2 (Version 2024.10) ^56^. Trimming parameters were selected that the average phred score across retained bases exceeded 30 for both 16S and ITS amplicons forward and reverse reads. Amplicon equencing variants (ASVs) were inferred using DADA2 Pipeline implemented in QIIME2 ^57^.

Taxonomic classification was performed using QIIME2 feature-classifier ^58^ with a custom rRNA/ITS operon reference database constructed from the genomes of the species used in this study. Chloroplast and low abundant ASVs (fewer than 50 total reads and present in more than four samples) were removed using phyloseq R package ^59^. ASV sequences that could not be classified on genus level were subsequently submitted to BLAST to enhance taxonomic resolution. Finally, relative abundances were calculated based on filtered ASV tables using phyloseq.

### Pseudomonas syringae CFU-determination from gnotobiotic in planta experiments

For bacterial quantification, plant material was homogenized in 10 mM MgCl₂ using zirconia beads. Plants were transferred to pre-weighed tubes containing 1 mL MgCl₂ and a mixture of 3-, 2-, 0.5-, and 0.1- mm zirconia beads, and fresh weight was determined. Samples were lysed using a Precellys homogenizer (four cycles at 6300 rpm for 30 seconds with 15 second pauses). Homogenates were serially diluted in MgCl₂ in 96-well plates, and aliquots were plated onto NB agar supplemented with rifampicin and cycloheximide. Rifampicin was used to selectively recover *Pseudomonas syringae*, which is resistant to the antibiotic, whereas other bacterial community members are not, and cycloheximide was included to suppress the growth of yeasts and filamentous fungi. Plates were incubated at room temperature, and colonies for each dilution were counted after 24 and 48 hours to calculate colony-forming units per millilitre.

## Data availability

All data needed to evaluate the conclusions in the paper are present in the paper or the Supplementary Materials. Raw amplicon sequencing data have been deposited in the NCBI Sequence Read Archive (SRA) under BioProject PRJNA1413978. Whole-genome sequencing data and the corresponding genome assemblies are available in the European Nucleotide Archive (ENA) under project accession PRJEB106563. Metabolomics raw data (previously centroided mzML MS/MS files) are deposited on Zenodo under DOI 10.5281/zenodo.18199851. Pangenome analysis datasets and all analysis code generated in this study are publicly available in the same Zenodo repository.

## Acknowledgments

We thank Libera Lo Presti for proofreading and helpful discussions. We also thank Abzer K. Pakkir Shah and Monja Schmid for their valuable suggestions. We thank the Weigel lab at the Max Planck Institute for Biology in Tübingen for kindly providing *Pseudomonas viridiflava* strains. We acknowledge support from the NGS Competence Center Tübingen (NCCT) and the Quantitative Biology Center (QBiC), University of Tübingen, Germany. ChatGPT-5 (OpenAI) was used for proofreading and text editing. All original content was authored by the authors, who take full responsibility for the manuscript.

## Author information

P.S., E.K., and D.P. conceptualized and designed the study and acquired funding. P.S. administered the project. P.S., L.B., C.B., M.N.D., A.P.L., S.F., J.B., K.S.L., M.M., V.C., D.R., N.Z., A.T.A., C.M.S., E.K., and D.P. performed the experiments and data analyses. P.S., E.K., and D.P. interpreted the data and wrote the manuscript. All authors revised the manuscript and approved the final version for submission.

## Extended data figures and tables

**Extended Data Fig. 1.**
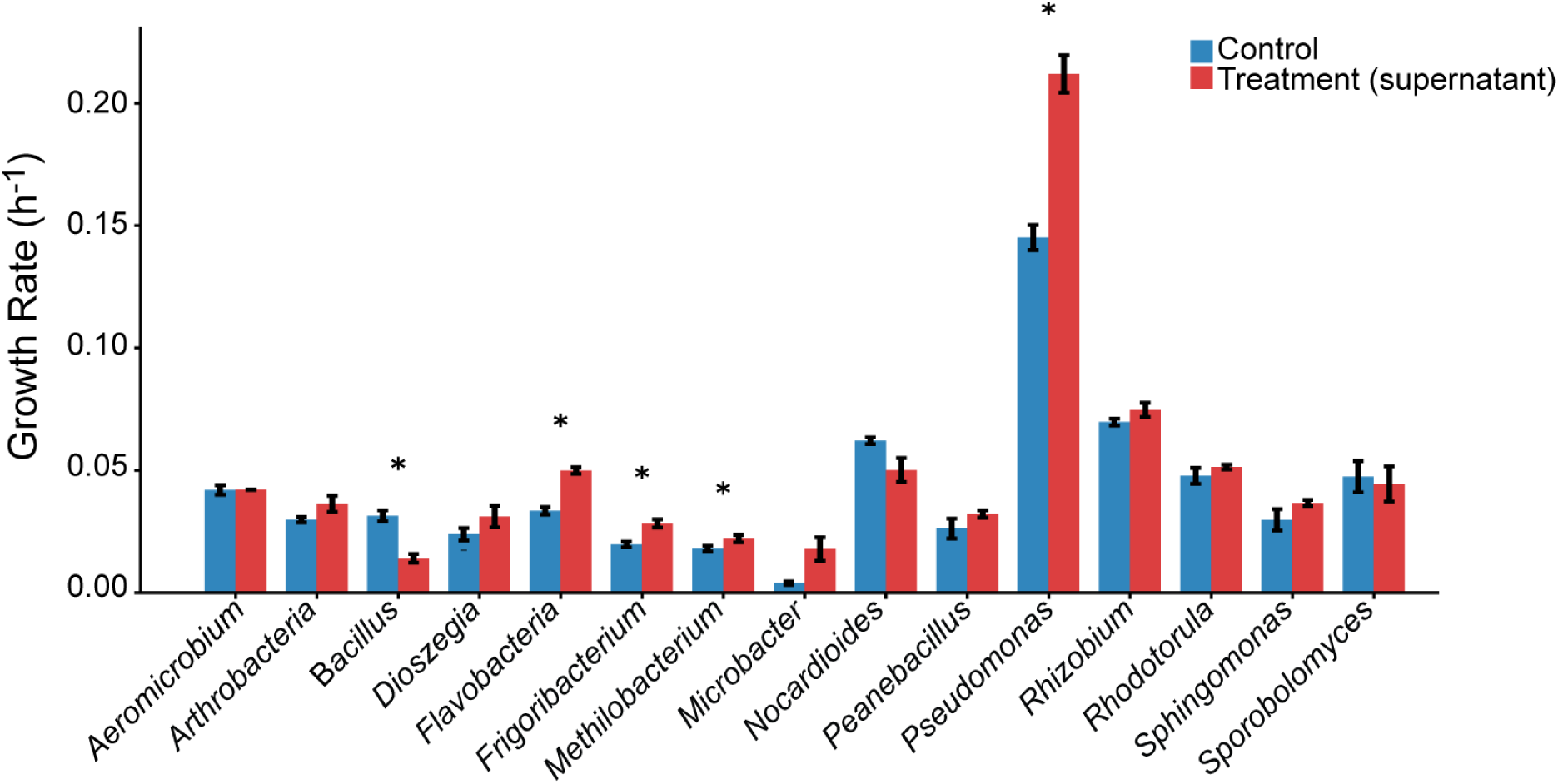
*Pseudomonas* growth is promoted by *Rhodotorula* supernatant. Bars show exponential growth rates of the 15 microbial strains comprising the phyllosphere core synthetic community (SynCom). Blue bars indicate growth in 100% fresh MM9 medium (control), whereas red bars indicate growth in a 50:50 (v/v) mixture of fresh MM9 and cell-free supernatant from a *Rhodotorula* preculture grown under iron-limited MM9 conditions, which stimulate RA production. Error bars denote standard deviation across replicate wells. Asterisks indicate significant differences between control and supernatant conditions (Welch’s t-test, *P* < 0.05).

**Extended Data Fig. 2.**
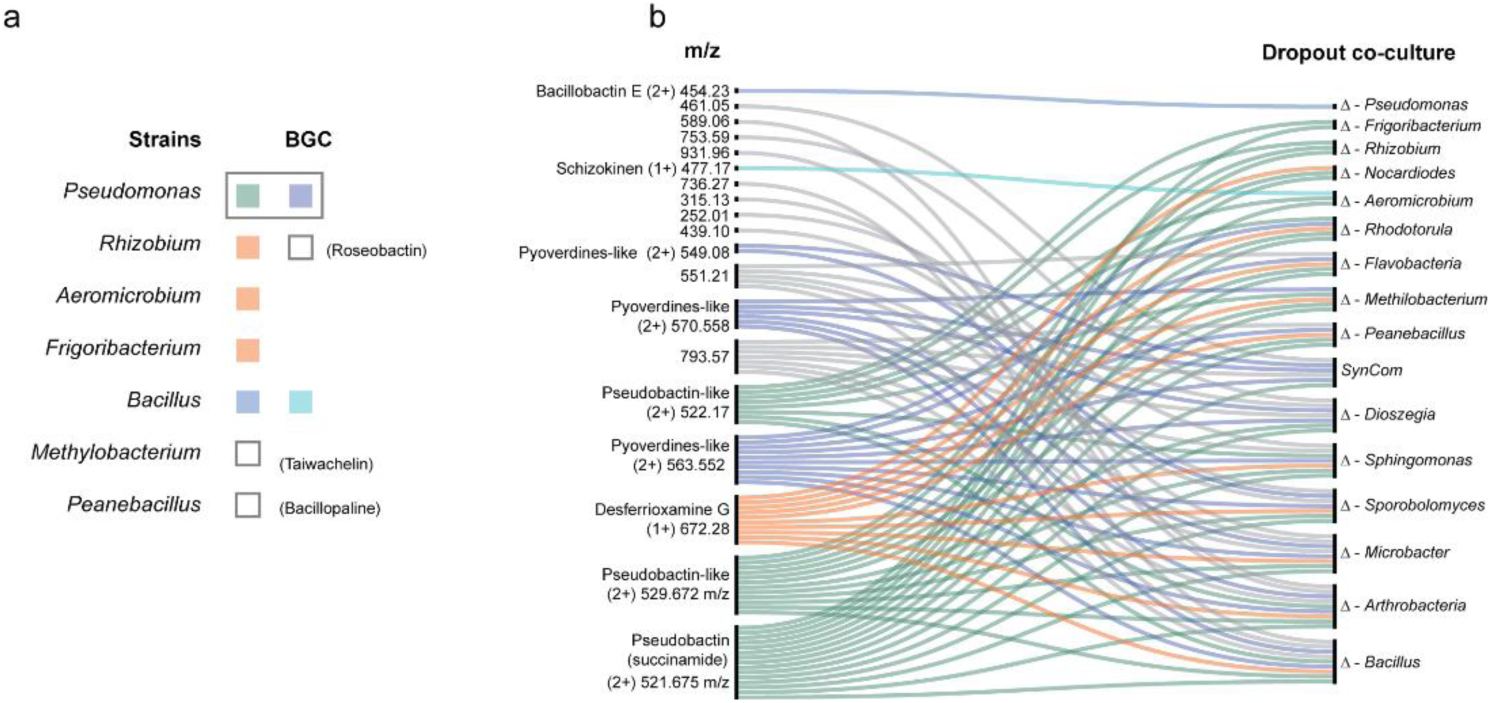
Mapping siderophore production in SynCom dropout co-cultures. **a,** SynCom strains harboring biosynthetic gene clusters (BGCs) for siderophore production, annotated using antiSMASH. Strains are color-coded according to the corresponding siderophore masses reported in the alluvial plot, which depicts relationships between community members and their siderophore production profiles. **b,** Alluvial plot summarizing MassQL results, showing detected *m/z* values corresponding to query-targeted siderophore masses in both singly and doubly charged states. *m/z* values were extracted directly from raw mass spectrometry data and reflect the microbial cultures in which these features were detected. Siderophore biosynthesis, as with other community metabolites, is modulated by microbial interactions. In the SynCom, specific dropout co-culture conditions alter the capacity of individual community members to produce siderophores. Selected masses were validated by native metabolomics and Ion Identity Molecular Networking (IIMN) analysis, including the siderophore pseudobactin (*m/z* 521.675), whose apo-form was not visible in the IIMN cluster due to its high iron-chelating capacity, and desferrioxamine G (*m/z* 563.552).

**Extended Data Fig. 3.**
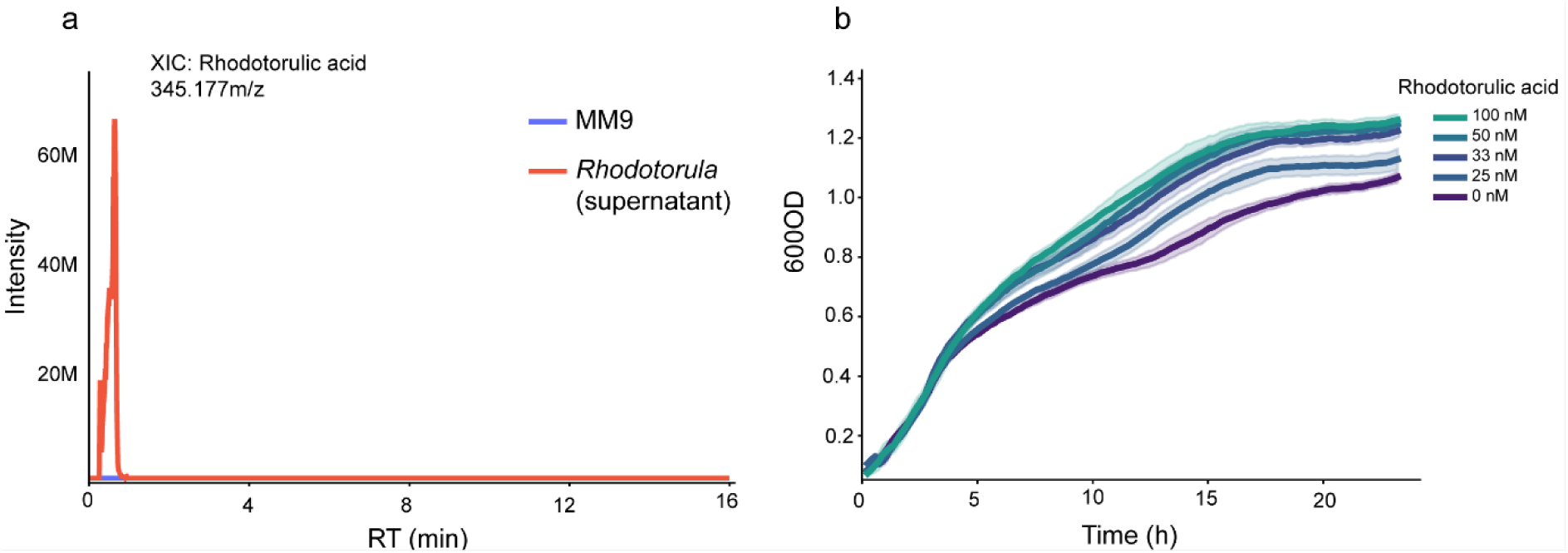
Growth of *Pseudomonas koreensis* across decreasing rhodotorulic acid concentrations. **a,** Extracted ion chromatogram (XIC) of rhodotorulic acid (345.177m/z) detected in the exometabolome of *Rhodotorula* culture supernatant. MM9 minimal medium served as a negative control and showed no detectable rhodotorulic acid signal. **b,** Growth curves of the SynCom *Pseudomonas koreensis* strain in MM9 minimal medium supplemented with decreasing concentrations of purchased rhodotorulic acid siderophore. MM9 without siderophore served as the control. Rhodotorulic acid enhanced growth, with maximal stimulation at ∼100 nM; this effect progressively diminished at lower concentrations, consistent with concentration-dependent siderophore utilization for iron acquisition.

**Extended Data Fig. 4.**
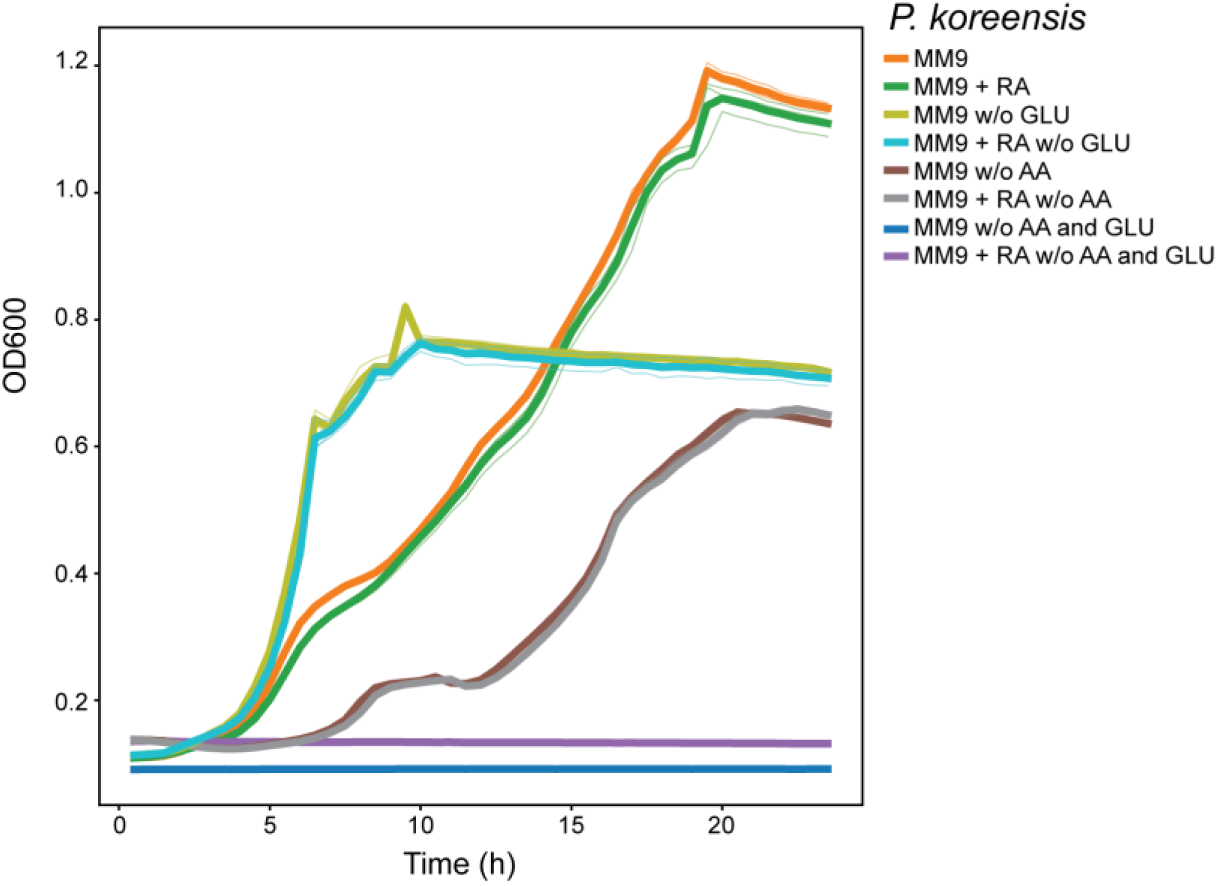
Growth of *Pseudomonas koreensis* under varying nutrient conditions and rhodotorulic acid supplementation. Growth curves of the SynCom *Pseudomonas koreensis* strain in MM9 minimal medium under the following conditions: standard MM9 ± rhodotorulic acid (50 µg mL⁻¹); MM9 lacking glucose ± rhodotorulic acid; MM9 lacking amino acids ± rhodotorulic acid; and MM9 lacking both glucose and amino acids ± rhodotorulic acid. Glucose depletion increased growth rate, whereas rhodotorulic acid supplementation did not support growth in carbon- or amino-acid–limited conditions, indicating it is not utilized as a carbon source.

**Extended Data Fig. 5.**
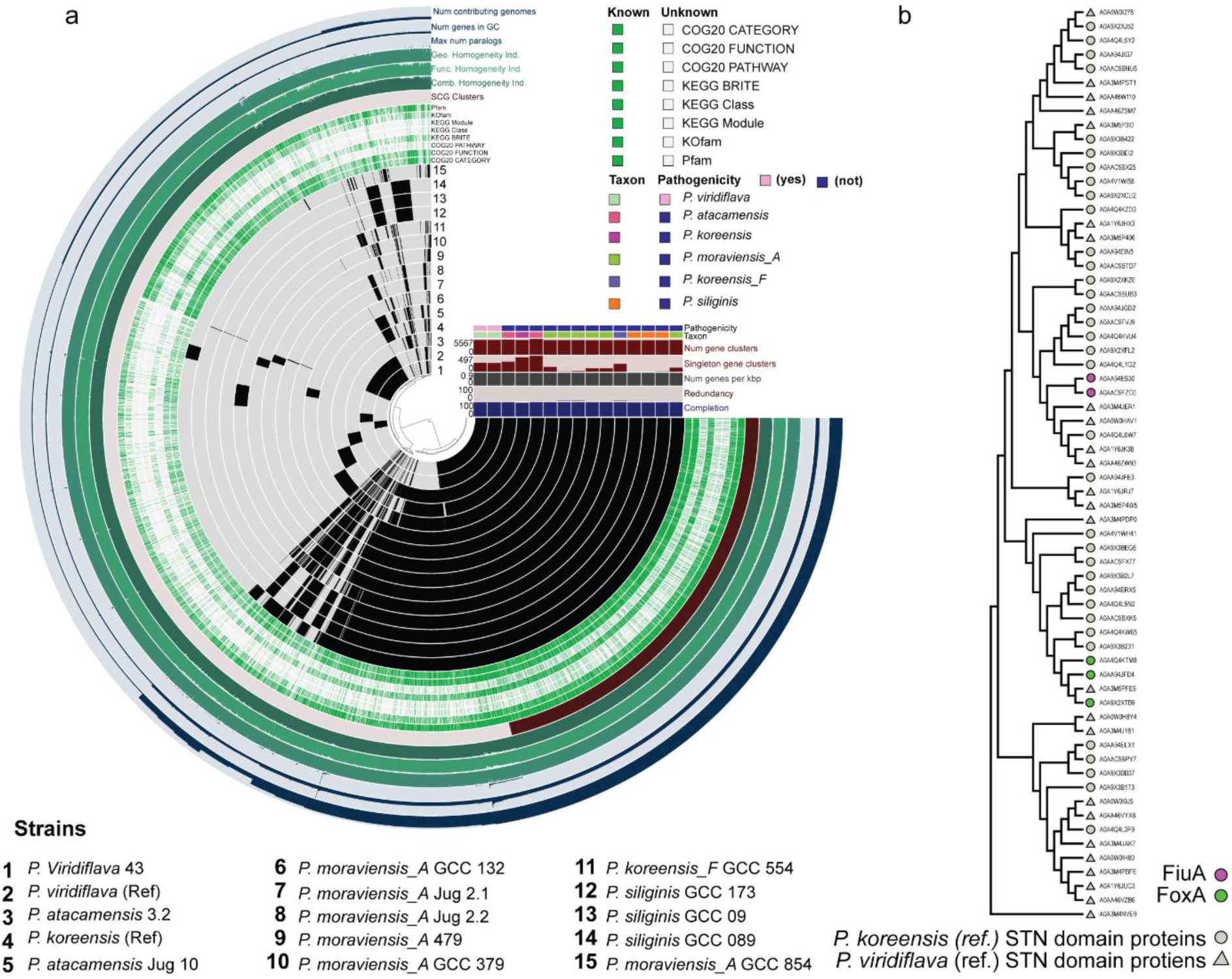
Whole-genome sequencing comparison of plant-isolated strains. **a,** Phylogenetic relationships of plant-associated *Pseudomonas* isolates. Eleven commensal strains clustered with the *P. koorensis* reference genome, whereas the weakly pathogenic *P. viridiflava* strain P43 grouped with the reference genome of the same species. **b,** Phylogeny of STN-domain proteins from *P. koorensis* and *P. viridiflava* reference genomes. Protein sequences were annotated using COG and PFAM domains and used to construct a Maximum Likelihood tree, with initial topology generated by Neighbor-Joining using pairwise JTT distances. The analysis included 65 amino acid sequences.

**Extended Data Fig. 6.**
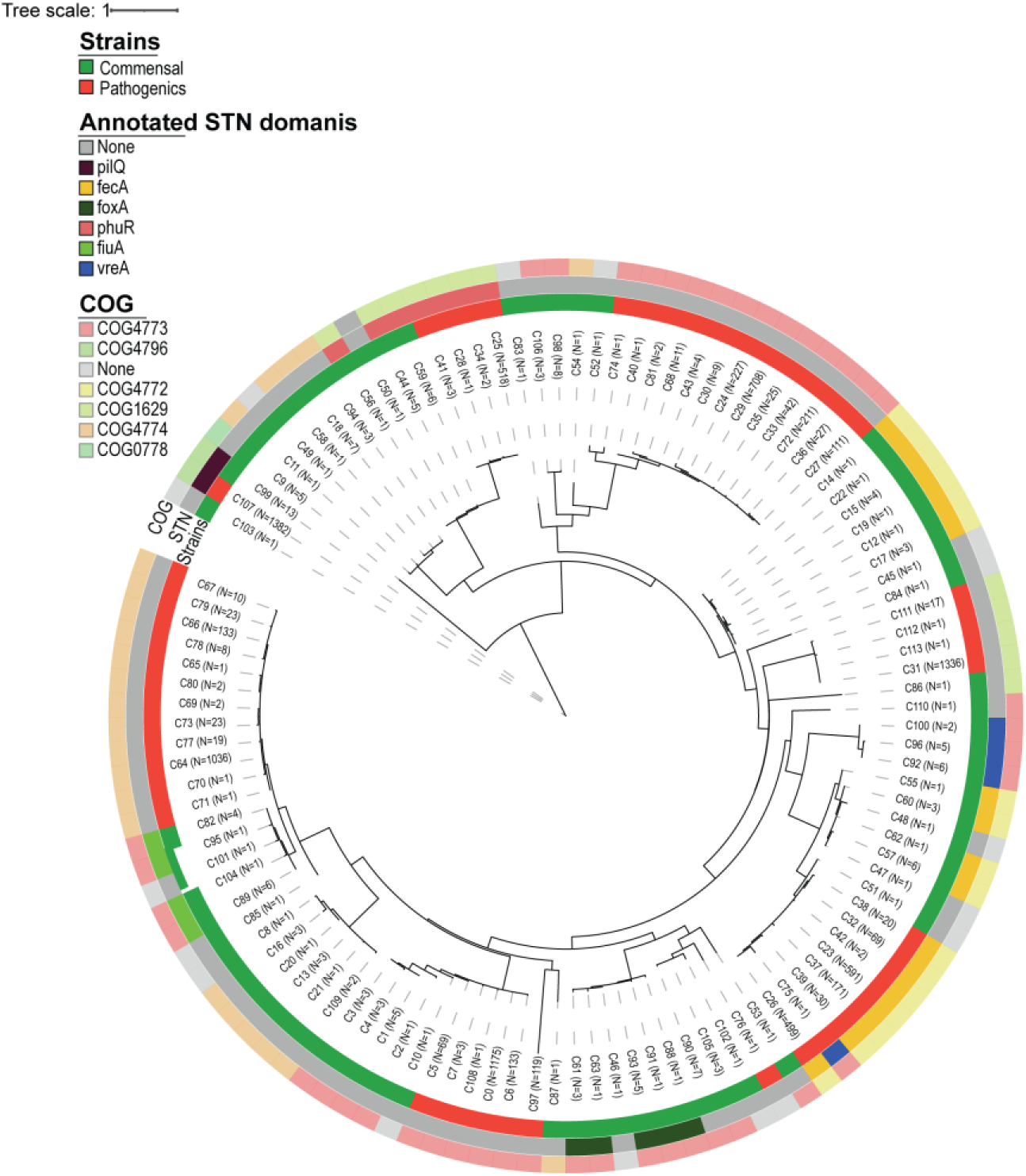
Phylogenetic analysis of TonB-box amino acid sequences. A clear divergence between commensal strains sequenced in this study and publicly available *Pseudomonas viridiflava* genomes. The TonB-dependent receptors fiuA and foxA are exclusively detected in the commensal strains.

**Extended Data Fig. 7.**
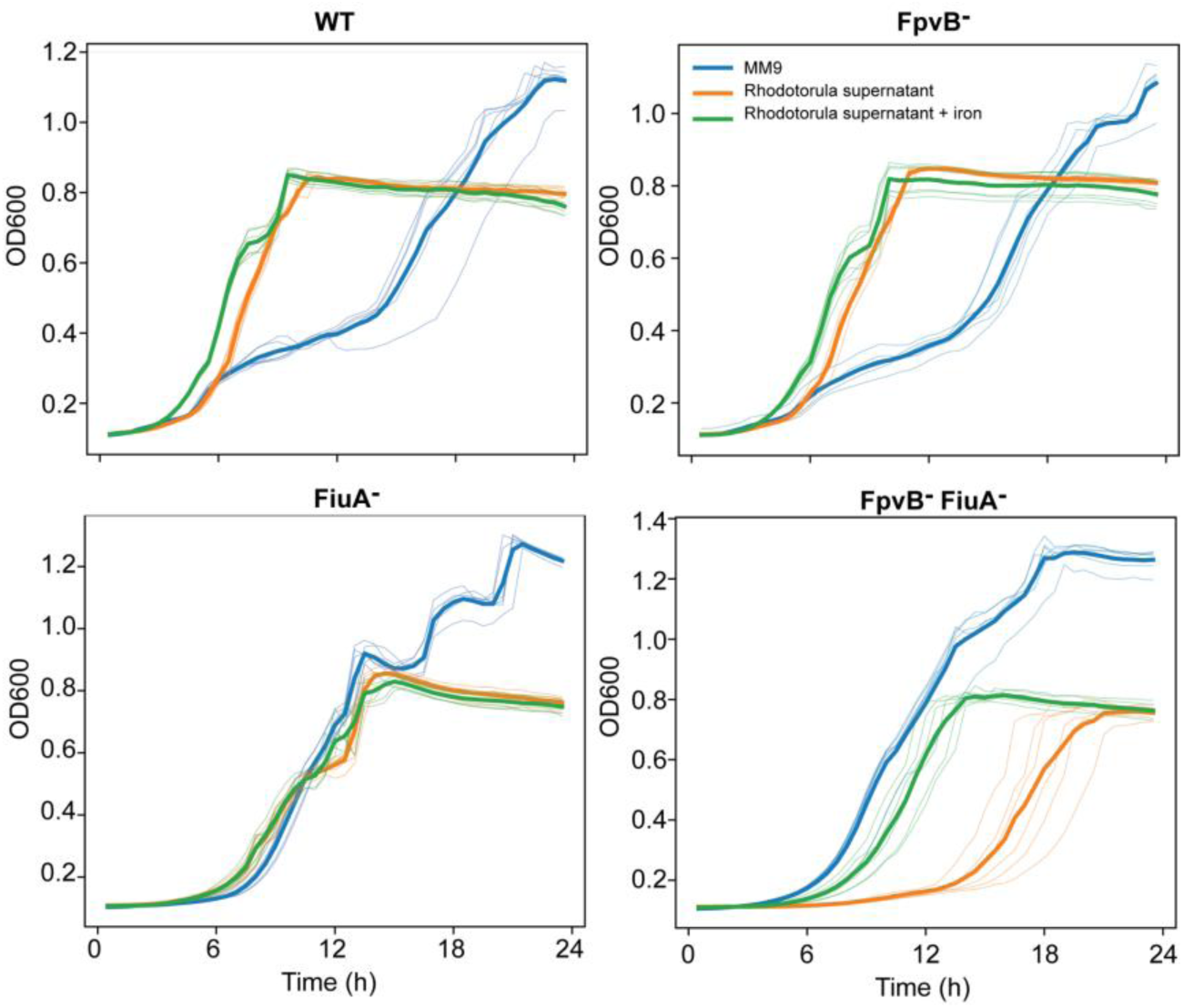
Iron limitation constrains the growth of *Pseudomonas koreensis* strains deficient in rhodotorulic acid uptake system. Growth dynamics of *Pseudomonas koreensis* SynCom mutants lacking TonB-dependent transporters in MM9 minimal medium and in *Rhodotorula* culture supernatant, including conditions in which the supernatant was supplemented with an excess of iron to saturate the RA present in the supernatant. Impaired growth in the supernatant results from the absence of rhodotorulic acid transport and is rescued by iron supplementation, indicating that iron availability is a key determinant of growth in strains deficient in rhodotorulic acid uptake.

**Extended Data Fig. 8.**
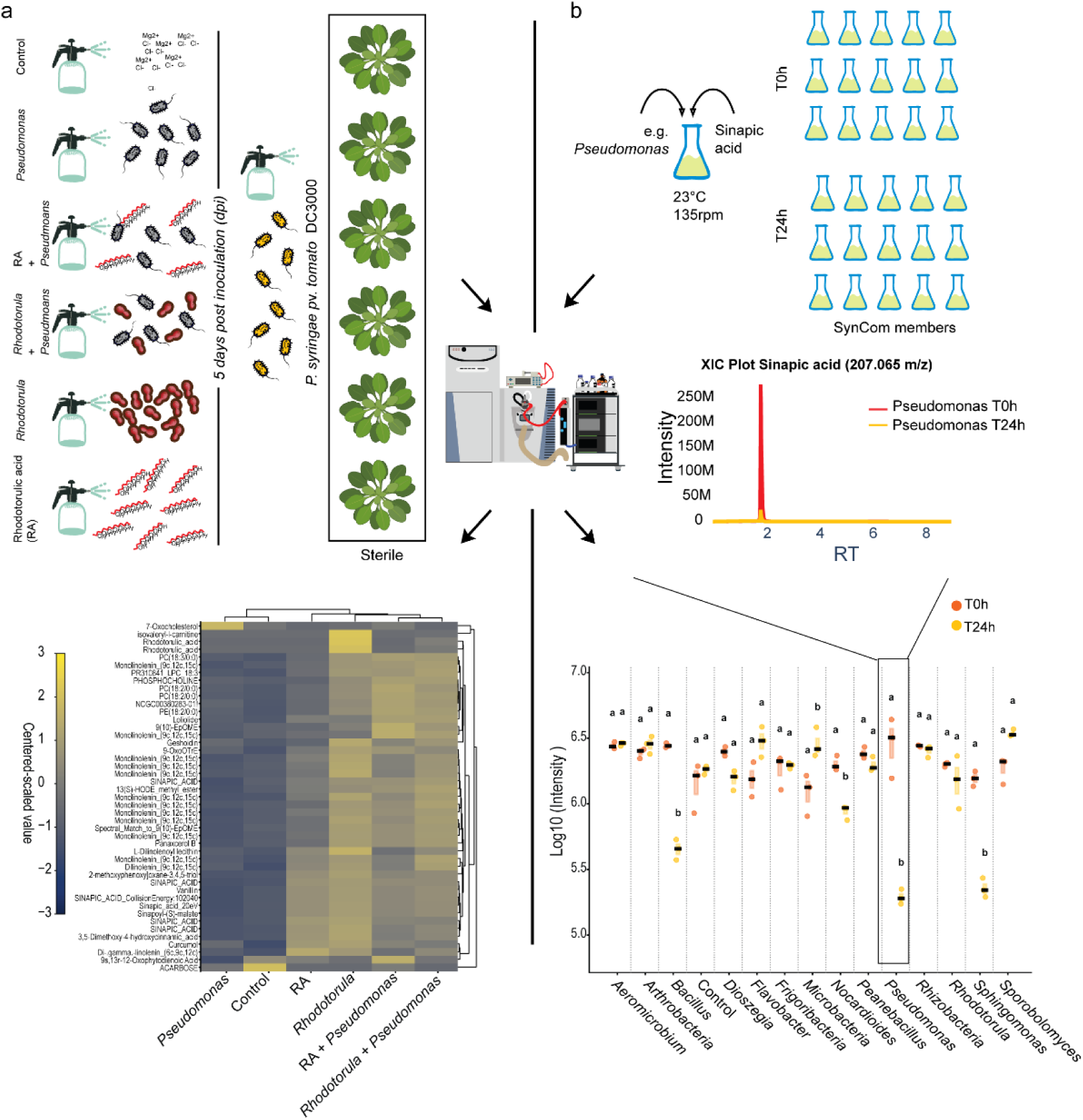
Rhodotorulic Acid promotes Phyllosphere through the Plant Metabolome. **a,** Gnotobiotic *Arabidopsis thaliana* plants were grown under sterile conditions and treated with Pseudomonas, *Rhodotorula* (O.2 600OD) rhodotorulic acid (30nM) alone, combination of microbial members and RA and untreated as control (MgCl2 10mM). Untargeted metabolomics followed by Feature-Based Molecular Networking (FBMN) revealed upregulation of sinapic acid and other phenolic acid related. **b,** To assess the impact of sinapic acid on the core synthetic microbial community (SynCom), individual strains were inoculated with the compound (100uM). Samples were collected at 0 and 24 hours to monitor microbial metabolic activity in the presence of sinapic acid. Untargeted metabolomics showed that several core SynCom members metabolized sinapic acid, as indicated by its depletion 24 hours after co-inoculation with the microbes.

**Extended Data Fig. 9.**
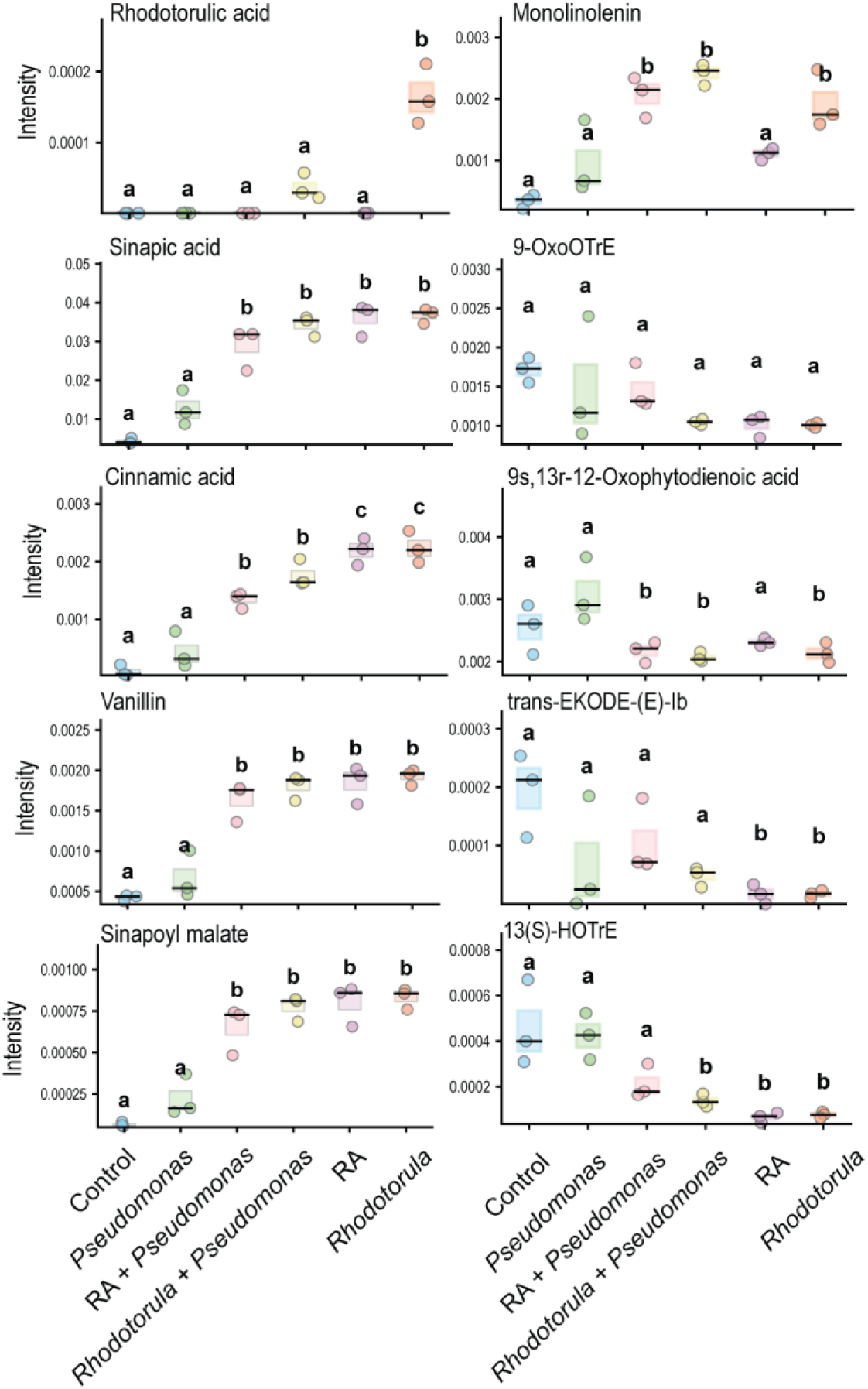
Differential regulation of lignin and jasmonic acid precursors during interkingdom interactions. Gnotobiotic plants were pre-inoculated with *Pseudomonas*, *Rhodotorula*, the siderophore RA, or their combinations prior to infection with *Pseudomonas syringae*. Lignin precursors accumulation (Sinapic acid, Cinnamic acid, Vanillin, Sinapoyl malate) were associated with depletion of jasmonic acid precursors (9-OxoOTrE, 9s,13r-12-Oxophytodienoic acid, Trans Ekode e-lb, 13(S)-HOTrE), particularly under Rhodotorula and RA treatment, consistent with reduced iron availability on leaf surfaces. One-way ANOVA (*P* < 0.05) followed by Tukey’s HSD test; different letters indicate significant differences (*P* < 0.05).

**Extended data Fig. 10.**
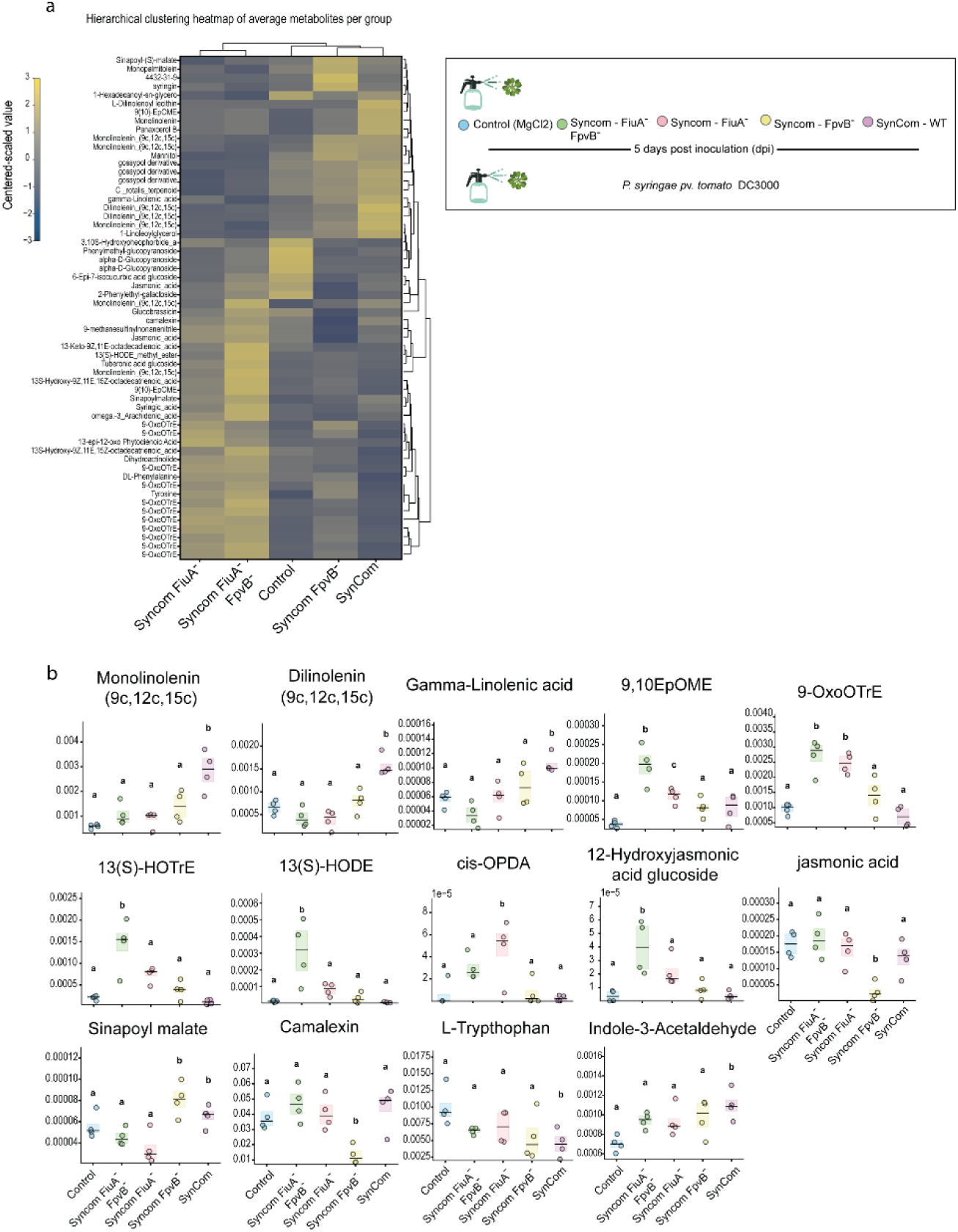
Non-targeted metabolomics of sterile plants exposed to SynCom variants containing substituted *Pseudomonas* strains. **a,** Sterile *Arabidopsis thaliana* plants were treated with SynCom subsets in which the commensal *Pseudomonas* member was replaced by knockout strains. After 5 days of community establishment, plants were inoculated with *Pseudomonas syringae* and harvested 7 days post-infection. Centered and scaled intensities of significantly regulated GNPS-annotated metabolites were averaged by treatment and subjected to hierarchical clustering (average linkage, Euclidean distance). The heatmap displays metabolites (rows) and treatments (columns) ordered by similarity; colors indicate relative abundance (blue to yellow, −3 to +3). Dendrograms depict hierarchical relationships among metabolites and treatments. **b,** Polyunsaturated fatty acids and jasmonic acid (JA) and auxin precursors were differentially regulated in plants colonized by the full community, partly recapitulating patterns observed upon inoculation with *Pseudomonas* and *Rhodotorula* alone. This trend was inverted in communities containing knockout strains, with increased JA precursors and reduced polyunsaturated fatty acids. Among phenolic, lignin-associated metabolites, only sinapoyl malate was differentially regulated, showing increased abundance in SynComs containing either the wild-type or the Fpvb mutant strain. One-way ANOVA (*P* < 0.05) followed by Tukey’s HSD test; different letters indicate significant differences (*P* < 0.05).

**Supplementary Table 1.** Summary of the Ion Identity Molecular Network (IIMN) edges, including Fe adducts. Edges connect metabolites (nodes) with iron adducts detected in the native metabolomics experiments in which iron salts were added to the system (Fig. 2F), including the siderophores rhodotorulic acid and desferrioxamine G.

| Name | scan1 | m/z | GNPS library | scan2 | m/z | GNPS library | EdgeAnnotation |
| --- | --- | --- | --- | --- | --- | --- | --- |
| 85917<br>(-)<br>85951 | 85917 | 518.32401 | PC(18:3/0:0);<br>[M+H]<sup>+</sup><br>C26H49N1O7P1 | 85951 | 286.6249 | #N/A | [M+Fe+2]<sup>+</sup>+2 [2M+2H]<sup>+</sup>+2<br>dm/z=231.69907 |
| 85884<br>(-)<br>85951 | 85884 | 1035.6385 | #N/A | 85951 | 286.6249 | #N/A | [M+Fe+2]<sup>+</sup>+2 [2M+H]<sup>+</sup><br>dm/z=749.01351 |
| 89758<br>(-)<br>89842 | 89758 | 480.30827 | Spectral Match to<br>1-(9Z-Octadecenyl)-<br>sn-glycero-3-phosphoethanolamine from<br>NIST14 | 89842 | 267.6173 | #N/A | [M+Fe+2]<sup>+</sup>+2 [M+H]<sup>+</sup><br>dm/z=212.69094 |
| 90415<br>(-)<br>90241 | 90415 | 288.64064 | #N/A | 90241 | 522.3553 | Spectral Match to 1-(9Z-Octadecenyl)-sn-glycero-3-phosphocholine from NIST14 | [M+Fe+2]<sup>+</sup>+2 [M+H]<sup>+</sup><br>dm/z=233.71462 |
| 104651<br>(-)<br>104701 | 104651 | 324.28943 | #N/A | 104701 | 378.2088 | #N/A | [M+Fe+2-H]<sup>+</sup>+ [M+H]<sup>+</sup><br>dm/z=53.91933 |
| 10623<br>(-)<br>11099 | 10623 | 689.34642 | #N/A | 11099 | 398.0882 | #N/A | [M+Fe+3-2H]<sup>+</sup>+ [2M+H]<sup>+</sup><br>dm/z=291.25819 |
| 10848<br>(-)<br>11099 | 10848 | 345.17671 | Rhodotorulic acid | 11099 | 398.0882 | #N/A | [M+Fe+3-2H]<sup>+</sup>+ [M+H]<sup>+</sup><br>dm/z=52.91153 |
| 32289<br>(-)<br>33426 | 32289 | 619.36629 | Desferrioxamine<br>G | 33426 | 336.6423 | #N/A | [M+Fe+3-H]<sup>+</sup>+2 [M+H]<sup>+</sup><br>dm/z=282.72401 |
| 36653<br>(-)<br>36511 | 36653 | 420.04145 | #N/A | 36511 | 786.1647 | FLAVIN ADENINE<br>DINUCLEOTIDE | [M+Fe+3-H]<sup>+</sup>+2 [M+H]<sup>+</sup><br>dm/z=366.12326 |
| 96862<br>(-)<br>97137 | 96862 | 610.36758 | #N/A | 97137 | 279.2317 | Spectral Match to 9(10)-<br>EpOME from NIST14 | [M+H]<sup>+</sup>+ [2M+Fe+3-2H]<sup>+</sup><br>dm/z=331.13592 |
| 9952 (-)<br>10848 | 9952 | 742.25815 | #N/A | 10848 | 345.1767 | Rhodotorulic acid | [M+H]<sup>+</sup>+ [2M+Fe+3-2H]<sup>+</sup><br>dm/z=397.08144 |
| 97068<br>(-)<br>96862 | 97068 | 261.22113 | #N/A | 96862 | 610.3676 | #N/A | [M-H2O+H]<sup>+</sup>+ [2M+Fe+3-2H]<sup>+</sup><br>dm/z=349.14645 |
| 104502<br>(-)<br>104361 | 104502 | 325.27343 | #N/A | 104361 | 396.1954 | #N/A | [M-H2O+H]<sup>+</sup>+ [M+Fe+3-2H]<sup>+</sup><br>dm/z=70.92199 |
| 104361<br>(-)<br>104523 | 104361 | 396.19542 | #N/A | 104523 | 342.3 | #N/A | [M-H2O+NH4]<sup>+</sup>+ [M+Fe+3-2H]<sup>+</sup><br>dm/z=53.89542 |

**Supplementary Table 2.** Whole-genome sequencing of commensal strains used in this study, with associated metadata including isolation site and genome features.

| Strain | Total<br>Read<br>Pairs<br>(M) | Total<br>Bases<br>(Gbp) | Total<br>Read<br>Pairs<br>(Trimmed)<br>(M) | Total<br>Bases<br>(Trimmed)<br>(Gbp) | Assembled<br>Size (Mbp) | Num<br>Contigs | N50<br>(kb) | GTDB Assignment |
| --- | --- | --- | --- | --- | --- | --- | --- | --- |
| wgs_S12036Nr1 | 8.08 | 2.41 | 7.94 | 2.37 | 6.69 | 637 | 105.7 | s_Pseudomonas_E_koreensis_A |
| wgs_S12036Nr2 | 8.10 | 2.42 | 7.96 | 2.37 | 5.92 | 229 | 247.5 | s_Pseudomonas_E_siliginis |
| wgs_S12036Nr3 | 8.10 | 2.41 | 7.95 | 2.37 | 5.94 | 281 | 112.3 | s_Pseudomonas_E_moraviensis_A |
| wgs_S12036Nr4 | 8.07 | 2.41 | 7.92 | 2.37 | 5.97 | 244 | 174.2 | s_Pseudomonas_E_moraviensis_A |
| wgs_S12036Nr5 | 8.08 | 2.41 | 7.93 | 2.38 | 6.04 | 207 | 249.7 | s_Pseudomonas_E_koreensis_F |
| wgs_S12036Nr6 | 8.12 | 2.42 | 7.98 | 2.38 | 5.92 | 249 | 233.7 | s_Pseudomonas_E_siliginis |
| wgs_S12036Nr7 | 8.11 | 2.42 | 8.03 | 2.39 | 6.03 | 228 | 101.7 | s_Pseudomonas_E_viridiflava |
| wgs_S12036Nr8 | 8.04 | 2.41 | 7.90 | 2.36 | 5.92 | 241 | 225.9 | s_Pseudomonas_E_siliginis |
| wgs_S12036Nr9 | 8.10 | 2.42 | 7.93 | 2.37 | 6.03 | 289 | 305.8 | s_Pseudomonas_E_moraviensis_A |
| wgs_S12036Nr10 | 8.11 | 2.42 | 7.97 | 2.38 | 6.05 | 249 | 196.2 | s_Pseudomonas_E_moraviensis_A |
| wgs_S12036Nr11 | 8.11 | 2.42 | 7.96 | 2.38 | 5.97 | 282 | 154.9 | s_Pseudomonas_E_moraviensis_A |
| wgs_S12036Nr12 | 8.06 | 2.41 | 7.92 | 2.36 | 5.97 | 278 | 164.4 | s_Pseudomonas_E_moraviensis_A |

